# The variegation of human brain vulnerability to rare genetic disorders and convergence with behaviorally defined disorders

**DOI:** 10.1101/2022.11.12.516252

**Authors:** Elizabeth Levitis, Siyuan Liu, Ethan T. Whitman, Allysa Warling, Erin Torres, Liv S. Clasen, François M. Lalonde, Joelle Sarlls, Daniel C. Alexander, Armin Raznahan

## Abstract

Diverse gene dosage disorders (GDDs) increase risk for psychiatric impairment, but characterization of GDD effects on the human brain has so far been piecemeal and lacked simultaneous analysis of multiple brain features across different GDDs. Here, through multimodal neuroimaging of 3 aneuploidy syndromes (XXY, XYY, trisomy 21), we reveal considerable diversity in cortical changes across GDDs and imaging-derived phenotypes (IDPs). This variegation of IDP change underlines the limitations of studying GDD effects unimodally. Integration across all IDP maps reveals highly distinct architectures of cortical change in each GDD, along with partial coalescence onto a common spatial axis of cortical vulnerability. This common axis shows strong alignment with shared cortical changes in behaviorally defined psychiatric disorders, and is enriched for specific molecular and cellular signatures - offering a high-priority target for future translational research.

## Introduction

Gene dosage disorders (GDDs) - including recurrent sub-chromosomal copy number variations and chromosomal aneuploidies - are individually rare but collectively common risks for neuropsychiatric impairment, and also provide naturally occurring models for genetic influences on human brain development. However, to date, most neuroimaging studies of altered brain organization in GDDs have considered at most only a few imaging derived phenotypes (IDPs) when comparing different GDDs^1–4^. The sparse sampling of brain changes in GDDs limits our biological understanding of each GDD in its own right and prevents us from determining the full extent of convergence vs. divergence in brain changes across different GDDs. In particular, any convergent effects of different GDDs on the brain may represent a biological substrate for the shared capacity of distinct GDDs to increase risk for a common set of behaviorally defined neuropsychiatric disorders (BDDs) such as autism spectrum disorder and attention deficit hyperactivity disorder^5^. To explore this potential, one needs to profile a wider gamut of IDPs in multiple GDDs, quantify shared and distinct effects of different GDDs on the brain, and test if any shared effects of GDDs align with brain changes seen in BDDs.

Here, we detail the regional effects of multiple GDDs on multiple IDPs through multimodal neuroimaging of the three most common human aneuploidy syndromes — Klinefelter syndrome (47, XXY), 47, XYY, and Down syndrome (trisomy 21, T21). We focus on these three particular GDDs because: (i) they are all trisomies involving supernumerary dosage of large and distinct gene sets, (ii) they have each been robustly associated with large effect size changes in brain structure and function (XXY^9^, XYY^6^, and T21^7,8^), and (iii) they are important from a medical perspective given their population prevalence (1:576 males XXY, 1:851 males XYY, 1:592 births T21^9^) and capacity to increase risk for neuropsychiatric and cognitive impairment^10,5^.

Our study examines 15 different IDPs in each GDD from T1-weighted structural magnetic resonance imaging (sMRI), diffusion weighted imaging (DWI) and resting state functional MRI (rsFMRI). This unique dataset enables us to advance understanding of genetic effects on the human brain in three key directions. First, by generating 15 IDP change (ΔIDP) maps for each GDD group we substantially expand upon the set of brain features that have been studied to date in human aneuploidies and thereby gain a more complete view of regional brain vulnerability in each disorder. Second, by comparing all 45 ΔIDP maps (15 in each GDD) to one another, we directly test competing mechanistic models for the mapping of genetic effects on cortical organization. For example, GDDs may all impact the same set of IDPs but in different cortical regions, or they may impact the same cortical regions but manifest in different IDPs. Third, we combine information across all IDPs to define and characterize the principal spatial component of cortical change in each GDD - as a step towards testing for a potentially shared backbone of multimodal change that is generalizable across multiple GDDs. Finally, we assess the extent to which shared effects of different GDDs on the brain align with brain changes shared among the many diverse BDDs that are seen at elevated rates in GDDs.

## Results

### Mapping alterations in cortical brain structure and function across three chromosomal aneuploidies

We assembled sMRI, DWI, and rsFMRI data from three age-matched case-control cohorts spanning XXY, (total n = 191, 92 controls), XYY (total n=81, 47 controls), and T21 syndromes (total n=69, 41 controls). In each of the three case-control cohorts, we computed 15 aneuploidy-specific ΔIDP maps through the workflow depicted in **Figure 1** (see **Methods**). These maps represent the effect of having an additional X, Y, or 21st chromosome on the 15 IDPs at each of 344 cortical gray matter ROIs defined using the Human Connectome Project multimodal parcellation^11^. The IDPs examined were: cortical thickness (CT, thickness), surface area (SA, area), gray matter volume (GMV, volume), mean curvature (MeanCurv, meancurv), gaussian curvature (GausCurv), intrinsic curvature index (CurvInd), and folding index (FoldInd), fractional anisotropy (FA), geodesic anisotropy (GA), mean diffusivity (MD), radial diffusivity (RD), axial diffusivity (AD), tensor mode (TM; mode), regional homogeneity (ReHo), and amplitude of low frequency fluctuations (ALFF). See **Table S2** for effect sizes and FDR-corrected q-values computed with and without TTV correction.

**Figure 1.**
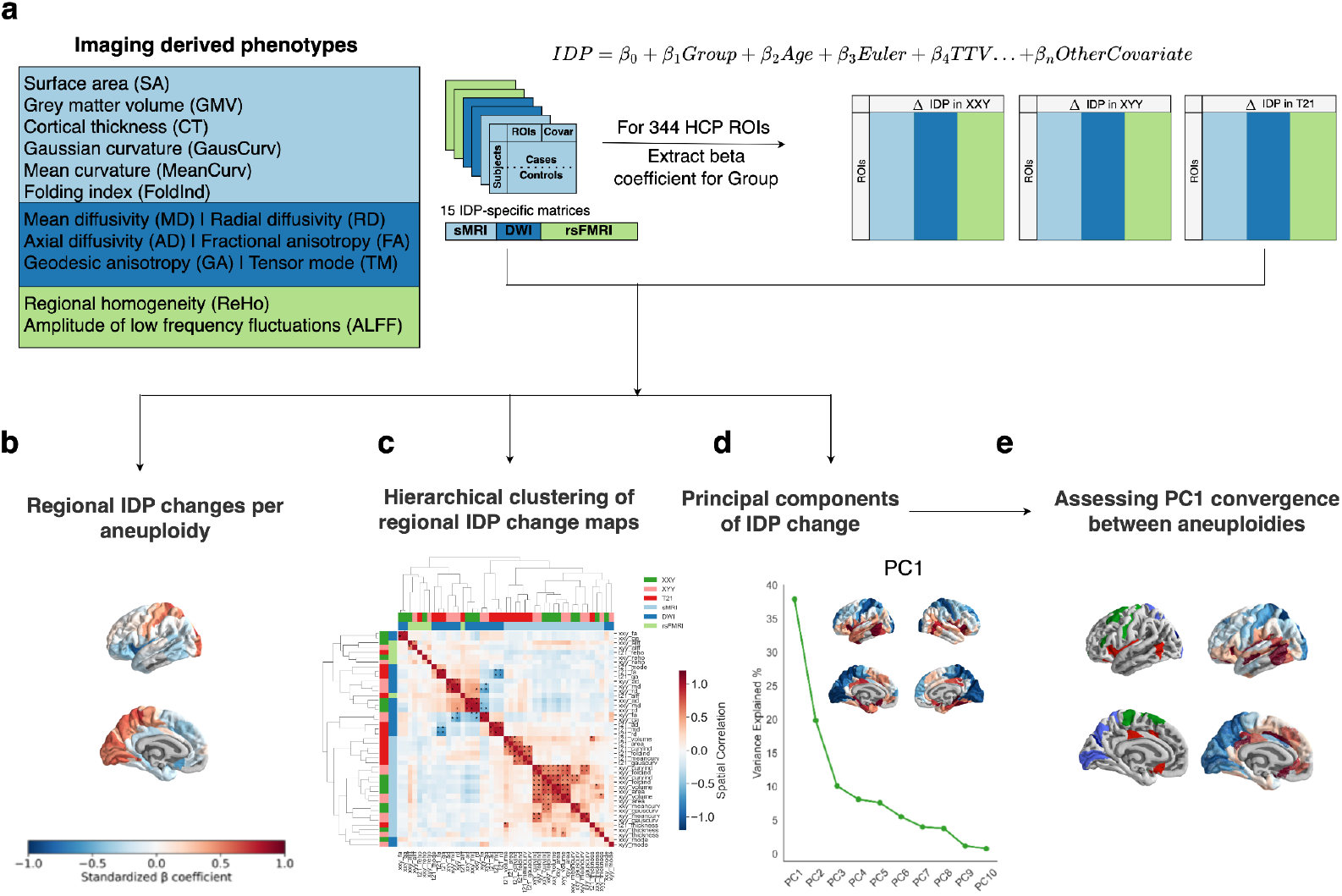
Analytical workflow. (**a**) On the left, we list the phenotypes analyzed for each modality. For each aneuploidy separately, we model each standardized phenotype as a function of group, age, euler number, total tissue volume, and aneuploidy + modality specific covariates. The group effect is extracted for each ROI and phenotype to yield a 344 ROI by 15 Δ IDP matrix per disorder. We (**b**) visualize the individual disorder-ΔIDP maps, (**c**) perform hierarchical clustering on a cross-disorder concatenated 344 by 45 matrix, (**d**) apply principal component analysis to each disorder’s ROI by ΔIDP matrix separately to identify patterns of multimodal change, and (**e**) assess PC1 convergence across aneuploidies continuously and discretely.

### Expanding the neuroimaging phenotypes of aneuploidy syndromes

We first examined individual ΔIDP maps in each aneuploidy to (i) verify that our findings replicated those of prior reports for those IDPs which have been previously studied (mainly for GMV, CT, and SA measures given the morphometric focus of past work), and (ii) assess the impact of each aneuploidy on the other IDPs that we examine for the first time (ReHo, ALFF, TM, FA, GA, MD, RD, and AD). For each aneuploidy, ΔIDP maps were visualized as un-thresholded effect size maps (**Figure 2a** left hemisphere, **Fig S2a** both hemispheres) and thresholded maps after FDR correction for multiple comparisons across regions (**Fig S2b** both hemispheres).

**Figure 2.**
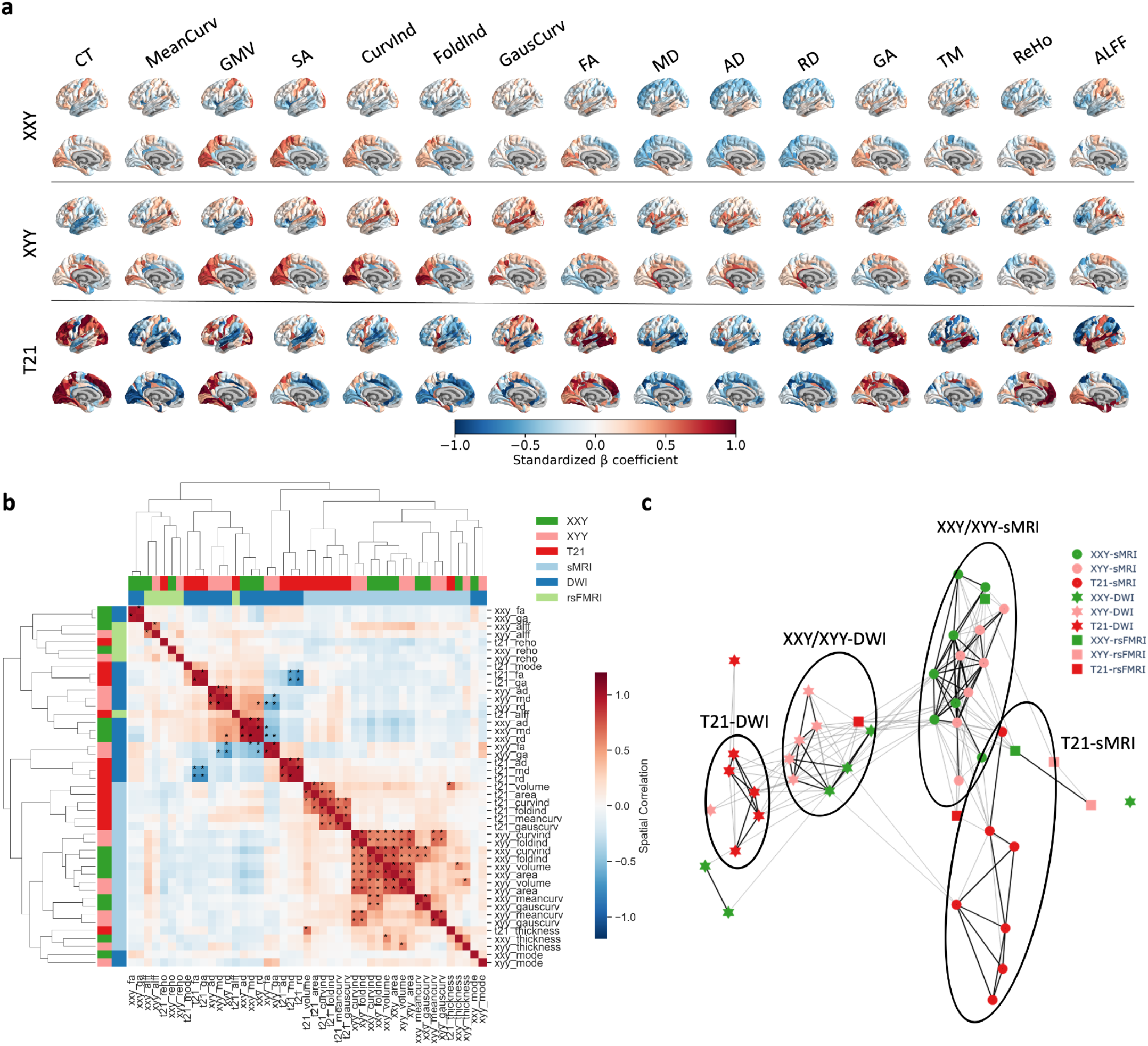
The spatial patterning of cortical change for 15 imaging-derived phenotypes (IDPs) in three aneuploidies. (**a**) Un-thresholded effect size maps of IDP change (i.e., “ΔIDP maps” for each aneuploidy (left hemisphere only, see **Fig S2a** for right hemisphere). Positive beta values reflect increases in cases relative to controls for a particular IDP of interest whereas negative beta values correspond to decreases in cases relative to controls. (**b**) Hierarchical clustering matrix of all 45 ΔIDP maps. Asterisks denote a significant spatial correlation between maps that survives correction using a strict null based on random spatial rotation (“spinning”) of maps. (**c**) A force-directed graphical representation of the relationship amongst all ΔIDP maps. Nodes represent ΔIDP maps, with color encoding the aneuploidy of origin and shape encoding imaging modality (squares - rsFMRI, circles - T1w, stars - DTI). Edges represent the spatial correlation between ΔIDP maps, edge width encoding the absolute Pearson r between maps across cortical ROIs and black edges being those that are statistically significant. Ellipses have been added to show the separation between T21-DWI nodes, XXY/XYY-DWI nodes, XXY/XYY-sMRI nodes, and T21-sMRI nodes from left to right.

For all three aneuploidies, our findings for GMV, SA and CT replicated those of prior reports^1,7^. For XXY and XYY we observed GMV and SA changes in bilateral frontotemporal (reductions) and occipitoparietal cortices (increases) as well as CT changes in temporoinsular (reductions) and somatosensory cortices (increases). ForT21 we observed occipotoparietal and somatosensory increases in GMV that were recapitulated by changes in CT while frontotemporal reductions in GMV were recapitulated by changes in SA. By newly detailing aneuploidy effects on sMRI-derived measures of cortical curvature we found widespread changes in cortical folding which echoed SA changes in each aneuploidy - suggesting that atypical gyrification is likely a key mediator of genetic effects on SA.

Examining effect size maps for DWI and rsFMRI derived IDPs enabled us to further expand understanding of regional cortical vulnerability to aneuploidy beyond the more commonly-studied morphometric features captured by sMRI. For instance, across the DTI-derived IDPs, FA alterations followed a rostrocaudal gradient in XYY (decreasing rostrally, increasing caudally), and a ventrodorsal gradient in XXY (decreasing dorsally, increasing ventrally), whereas FA showed sub-threshold global increases in T21. Diffusivity measures also showed regionally patterned alterations that varied between aneuploidies and differed from FA changes. Changes in rsFMRI-derived IDPs were also regionally specific in a manner that varied for ALFF and ReHo as well as across the aneuploidies. For example, ALFF was decreased in the inferior frontal cortex of both XXY and XYY while in XXY it was also decreased in the medial prefrontal cortices as well as in the anterior and posterior cingulate. In contrast ALFF showed large effect size parietofrontal decreases and lateralized insular increases in T21. Of note, the robust effect size of T21 for most sMRI-derived IDPs (**Fig 2a**) combined with the lack of statistically-significant effects for DWI- and rsFMRI-derived metrics is consistent with the reduced sample size for these two modalities relative to that for sMRI in T21 - a phenomenon which also applies to XYY and XXY cohorts and thus encourages cross-IDP visual comparisons using effect size maps (**Fig 2a**) rather than thresholded maps alone (**Fig S2b**). Nevertheless, qualitative comparison of both unthresholded (**Fig 2a**) and thresholded (**Fig S2b**) ΔIDP maps indicates that inclusion of DWI- and rsFMRI-derived phenotypes identified additional regional effects of aneuploidy on the brain that were not detected using only sMRI-derived phenotypes.

These results substantially advance our understanding of regional cortical involvement in XYY, XXY and T21 by extending the phenotypic depth with which cortical change is characterized. The diversity of regional involvement across different IDPs in each aneuploidy emphasizes the importance of examining multiple cortical features in parallel when studying GDDs. We observe extensive regional cortical involvement with unique spatial distributions when considering IDPs besides the most commonly studied morphometric features derived from sMRI (GMV, CT and SA).

### Defining organizing-principles of cortical change across IDPs, regions, and aneuploidies

To quantitatively compare cortical changes for different IDPs and aneuploidies, we computed cross-ROI spatial correlations across all 45 ΔIDP maps (15 maps for each of 3 aneuploidies). Hierarchical clustering of the resulting 45*45 symmetric matrix revealed that similarities between ΔIDP maps were most strongly organized by imaging modality, and then by aneuploidy type (**Fig 2b**). For example, when the correlation matrix is divided into three clusters, the ΔIDP maps were grouped into a large cluster of predominantly sMRI-based ΔIDP maps (e.g., CT, SA, GMV, FoldInd, GausCurv, MeanCurv) from all 3 aneuploidies. The remaining two clusters were mostly made up of DWI- and rsFMRI-based features - one dominated by ΔIDP maps for diffusivity, anisotropy, ReHo, and ALFF measures, and the other including T21 diffusivity and XYY anisotropy ΔIDP maps. The correlational structure within each of these clusters appeared to be primarily (but not exclusively) organized by aneuploidy type with subclusters indicating that the spatial pattern of cortical changes in T21 differs from that in XXY and XYY. To provide a more constrained view of these relationships, we thresholded the matrix of correlations between ΔIDP maps to retain only those that were statistically significant after implementation of a spatial permutation test and correction for multiple comparisons^12,13^. Seventy-five of all 990 unique inter-ΔIDP map correlations met this strict criteria (asterisked in **Figure 2b**), and we visualized these connections between ΔIDP maps as a force-directed graph where nodes are ΔIDP maps and edges encode the strength of spatial similarity between pairs of ΔIDP maps (**Fig 2c**). Reinforcing the patterns seen on hierarchical clustering of the ΔIDP correlation matrix, the clusters in this graph were primarily distinguished by imaging modality, and nodes within each cluster were organized by aneuploidy type. The strongest edges were between changes in anisotropy measures for each disorder, diffusivity measures for each disorder, volume and area for each disorder, and morphometric measures between XXY and XYY, and ALFF between XXY and XYY.

Thus, an agnostic comparison of cortical change maps spanning different phenotypes and aneuploidies points to overarching principles organizing regional cortical vulnerability in GDDs. Specifically, the dominant source of variation in the spatial pattern of cortical change in aneuploidy appears to be the imaging modality with which cortical change is being measured, and an important secondary source of variation is the specific aneuploidy being considered. Thus, the degree of convergence between the spatial pattern of cortical vulnerability to different GDDs varies depending on the specific IDPs and GDDs being considered.

### Defining and characterizing principal components of multimodal cortical change within and across aneuploidies

We next used principal component analysis (PCA) to provide an efficient summary representation of multimodal change for each aneuploidy - defined by ROI scores for the first principal component (PC1) of the aneuploidy’s 344*15 ΔIDP matrix and associated feature loadings for this PC. The three PC1 maps explained 38%, 31%, and 25% of the variance in XXY, XYY, and T21 ROI*ΔIDP matrices respectively (**Fig 3a**). The spatial distribution of regional PC1 scores showed some qualitative similarities, with all aneuploidy PC1 maps having high scores in perisylvian cortices and low scores in medial occipital regions (**Fig 3b**). Quantitatively, the cross-ROI correlation in PC1 scores varied substantially between aneuploidy pairs, being most similar between XXY and XYY (r=0.73) and least similar between T21 and the two sex chromosome aneuploidies (between XXY and T21: r=0.24, XYY and T21: r=0.30) (**Fig 3c**). Evident drivers of the low spatial correlation between PC1 in T21 and the two sex chromosome aneuploidies were the extreme negative occipitoparietal PC1 scores in XYY and XXY but not T21, and the presence of extreme negative medial temporal lobe PC1 scores in T21 but not XYY or XXY. The three aneuploidies also differed in the relative contribution of ΔIDP maps to PC1. Correlations in PC1 factor loadings were higher between the two sex chromosome aneuploidies (r=0.95) and lower when correlated with T21 (between XXY and T21: r=0.33, XYY and T21: 0.54; **Fig 3b, d**). In XXY and XYY, volumetric and diffusivity/ReHo measures tend to make counterpoised contributions to the spatial patterning of multimodal cortical change, while in T21, volumetric and anisotropy/ReHo measures have opposing contributions. We focus here on PC1 of multimodal change in each aneuploidy, but subsequent PCs up to a cumulative variance explained of 85% are shown on **Fig S5a, b.** These components of multimodal change were generally less consistent between aneuploidies than PC1 in their spatial distributions (**Fig S5c**); however, a given pattern of feature loadings could connect disparate PCs across aneuploidies (**Fig S5d**).

**Figure 3.**
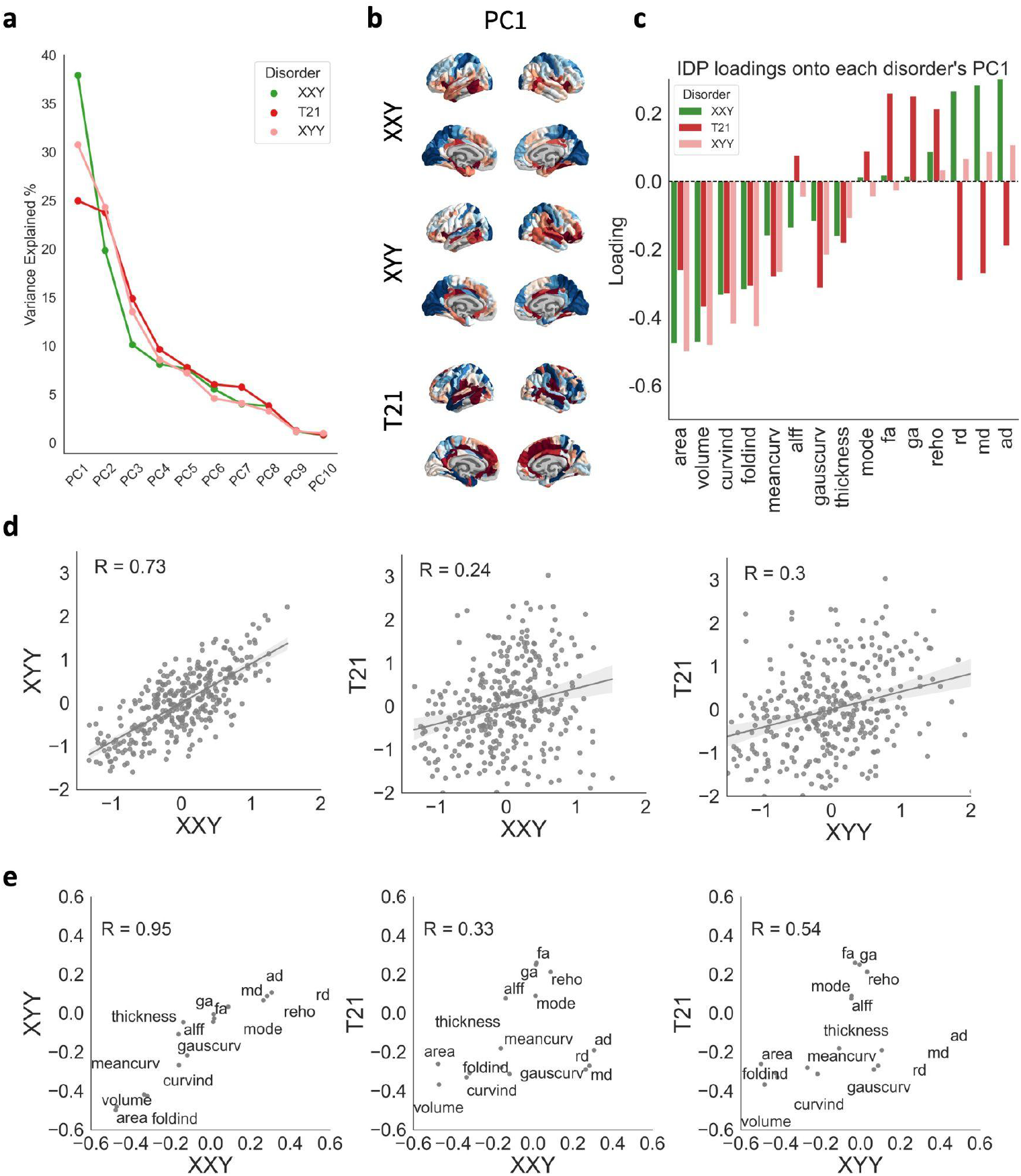
Defining and comparing the principal component of cortical change in each aneuploidy. **(a)** Proportion of variance explained by the top 10 principal components of each aneuploidy’s ROI by ΔIDP matrix. (**b**) ROI PC1 scores for each aneuploidy projected onto the surface of the brain. (**c**) IDP loadings for each aneuploidy’s PC1. (**d**) Correlation between regional PC1 scores for each pair of aneuploidies. (**e**) Correlation between PC1 IDP loading for each pair of aneuploidies.

Thus, in each aneuploidy approximately ⅓ of the spatial variation in 15 IDP changes across the cortex can be explained by a single principal spatial component which emphasizes morphometric (sMRI-based) and microstructural (DWI-based) measures. This spatial component is not identical between aneuploidies: it shows strong similarity between XXY and XYY in both regional scores and feature loadings, but weak to moderate similarity between the two sex chromosome aneuploidies and T21. Moreover, each aneuploidy shows several spatial components of multimodal cortical change beyond this first principal component, and these tend to be highly dissimilar across the aneuploidies. Taken together, these results indicate that GDDs differ greatly from each other in the full profile of their effects on human cortical anatomy, but there appears to be a primary spatial axis of anatomical change that is at least moderately similar between different GDDs.

### Convergent multimodal cortical changes across aneuploidies and links to convergence across behaviorally-defined disorders

Having defined PC1 of multimodal cortical change in each aneuploidy, we then assessed commonalities between these PC1 maps to identify potential signatures of shared cortical vulnerability between GDDs. To achieve this we averaged XYY, XXY, and T21 PC1 maps to provide a single summary map of human cortical vulnerability to aneuploidy. This cross-aneuploidy average PC1 map highlighted the insula and ventro-medial prefrontal cortex (high average PC1) as well as occipitoparietal cortices (low average PC1) as regions of core cortical vulnerability (**Fig 4a**). These regions of convergent vulnerability could also be recovered through a complementary conjunction analysis across extreme negative (0-10th decile PC1 scores, blue), extreme positive (90th-100th decile PC1 scores, red) and low (45th-55th decile PC1 scores, green) scores in the three aneuploidy-specific PC1 maps (**Methods, Fig 4b**). The conjunction of low PC1 scores across aneuploidies (green regions in **Fig 4b**) highlighted the temporoparietal junction and regions of the lateral prefrontal cortex as areas that could either have relative resilience or inconsistent vulnerability across aneuploidies.

**Figure 4.**
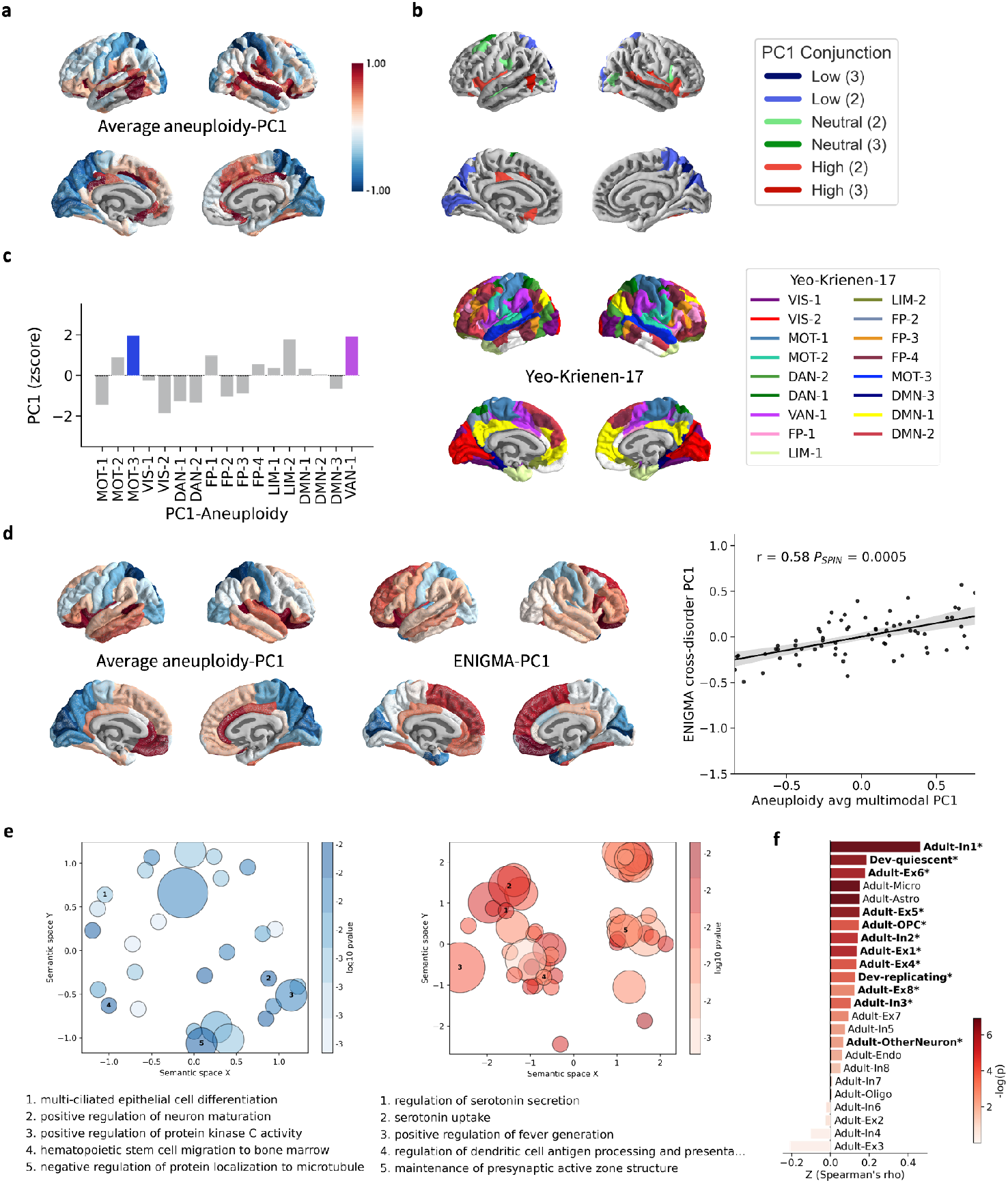
Defining and characterizing convergent regional vulnerability of the human cortex. (**a**) Average PC1 map (Aneuploidy-PC1) projected onto the cortical surface (in the HCP/Glasser atlas parcellation). (**b**) Conjunction map showing ROIs which sit at the top (red), middle (greens) and bottom (blues) deciles of PC1 scores across aneuploidy/GDD. (**c**) Enrichment of functional connectivity modules defined by Yeo-Krienen-17 (top cortical surfaces, mapped to the Glasser parcellation) for extreme PC1 scores within the Aneuploidy-PC1 map. (**d**) On the left, we show the average aneuploidy PC1 map re-parcellated using the Desikan Killiany atlas (see Methods). In the middle, we show the PC1 map generated from applying principal component analysis to case-control SA and CT maps from behaviorally defined psychiatric disorders in the ENIGMA dataset (ENIGMA-PC1). On the right, we show the spatial correlation between the Aneuploidy-PC1 map and ENIGMA-PC1 map. (**e**) We show the gene ontology (GO) biological processes’ GCEA results for the Aneuploidy-PC1 map. The two panels are color-coded in red and blue with red denoting categories that are significantly enriched for the high end of the PC1 map whereas blue denotes categories that are enriched for the low end of the PC1 map. Each circle represents a cluster of gene ontology (GO) categories, with the point size representing the number of categories in a cluster, point color represents the uncorrected p-values derived after fitting a standard distribution to the permutation-derived null category scores. Clusters are sorted by the cScorePheno scores. Only categories that are significant after FDR correction are included here. (**f**) Cell type enrichment analysis using bars that represent category scores, and the color reflects the negative base 10 logarithm of the uncorrected p-values, asterisks and bolding represent those cell types that survive FDR correction.

We next harnessed the cross-aneuploidy average PC1 map (**Fig 4a**) to ask if regions showing shared vulnerability to different GDDs are enriched for particular functional domains by using spin-based permutations to compare the average PC1 map with a canonical parcellations of the cortex into 17 different rsFMRI connectivity networks (Yeo-Krienen-17^14^). An omnibus test revealed non-random alignment between this parcellation and the average PC1 map (p_spin_ < 0.05: **Fig S7a**), and post hoc testing localized this alignment by showing a significant enrichment of high average PC1 scores in somatomotor (MOT-3) and ventral attention (VAN-1) subnetworks (**Fig 4c**). Of note, these networks were not consistently identified as showing statistically significant enrichment in individual analyses of each aneuploidy-specific PC1 map (**Fig S8b**), indicating that the shared effects of multiple GDDs on regional cortical organization highlight cortical systems that may not show concentrated cortical change in individual GDDs.

Finally, given that all three of the aneuploidies included in our study can increase risk for multiple behaviorally-defined neuropsychiatric disorders (BDDs)^5,10^ and that prior work has hinted at a shared spatial pattern of cortical vulnerability to multiple BDDs^15,16^, we tested for an overlap between shared GDD and shared BDD effects on human cortical anatomy. We turned to the ENIGMA dataset to obtain case-control maps for 21 SA and CT change maps from BDDs across the lifespan (see **Methods**), and we applied PCA to identify the first principal component (ENIGMA-PC1) of morphometric change across the maps. This principal component of cortical vulnerability in BDDs showed a moderate-strong correlation with the average aneuploidy PC1 map (R=0.58; p_spin_ = 0.0005; **Fig 4d**), indicating the existence of a core spatial axis of human cortical vulnerability to diverse genetically and behaviorally defined neuropsychiatric disorders. To better characterize the biological properties of this spatial axis we aligned the average PC1 map in aneuploidy to the Allen Human Brain Atlas (AHBA^17^)’ and used ensemble-based gene category enrichment analysis (GCEA) to quantify associations between the AHBA genes’ spatial expression and the average PC1 map in aneuploidy^17–19^ (see **Methods**). We first identified biological processes in which genes most associated with the Aneuploidy-PC1 phenotype were implicated, and found 371 biological process categories with significant enrichment. Enrichment scores (cScorePheno) were quantified using the mean of r-to-z-transformed phenotype-gene Spearman correlation coefficients for all genes in a category of interest. Forty biological process categories were enriched for genes negatively correlated with the average PC1 map (i.e., relatively high expression in regions with extreme negative PC1 scores), the most enriched categories being: “multi-ciliated epithelial cell differentiation” (cScorePheno = 0.21; *q*_FDR_ = 0) and “positive regulation of neuron maturation” (cScorePheno = 0.2; *q*_FDR_ = 0). In contrast, 331 categories were enriched for genes positively correlated with the average PC1 map (i.e., relatively high expression in regions with extreme positive PC1 scores) - the most enriched categories being related to serotonin and norepinephrine - “regulation of serotonin secretion” (GO:0014062; cScorePheno = 0.34; *q*_FDR_ = 0.047), “serotonin transport” (GO:0006837; cScorePheno = 0.33; *q*_FDR_ = 0.043), and “serotonin uptake” (GO:0051610; cScorePheno = 0.29; *q*_FDR_ = 0.043), “norepinephrine uptake” (“GO:0051620”; cScorePheno = 0.29; *q*_FDR_ = 0.047. These biological process category enrichments are visualized using GO-Figure!^20^ to cluster statistically significant GO-categories based on semantic similarity (**Fig 4e)**. Gene category enrichment analysis also revealed that genes positively correlated with the average PC1 map were significantly enriched for markers of 12 cell types (**Fig 4f**), with In1, a class of inhibitory interneurons, showing the highest enrichment (cScorePheno = 0.46; *q*_FDR_ = 0.008). Complete GCEA enrichment results for average PC1 are provided in **Table S4.**

Thus, our analysis of three aneuploidy syndromes suggests that while the full pattern of multimodal cortical change can be largely non-overlapping across different GDDs (**Fig S5a,b**), some modest commonalities can exist between GDDs in the first principal spatial component of multimodal cortical change (**Fig 3**). Averaging this component across the 3 aneuploidies studied suggests a potential core spatial axis of cortical vulnerability to GDDs which highlights the ventromedial prefrontal cortex and insula (strongly positive average PC1 scores) and occipito-parietal cortices (strongly negative PC1 scores) as poles of vulnerability to counterbalanced changes in cortical macro- and microstructure. Strikingly, this shared spatial axis of cortical vulnerability to different GDDs is closely aligned to the dominant spatial axis of cortical vulnerability to multiple BDDs, suggesting that diverse risk factors for neuropsychiatric illness can converge to modify a core set of brain regions.

### Stability of findings with and without controlling for total tissue volume

Previous studies have indicated that aneuploidies of chromosomes X, Y, and 21 significantly alter overall brain size (decreasing, increasing, and decreasing brain size, respectively), and that controlling for these global effects can modify maps of regional cortical anatomy change in each aneuploidy^1,7^. Thus, as a supplementary analysis, we evaluated the difference between the IDP change maps with and without correction for TTV (**Fig S2a, Fig S3a**). The maps looked most distinct for the morphometric measures in XXY and T21, reflecting the previously identified total decrease in brain volume for both aneuploidies. Spatial correlations between each TTV-corrected IDP change map and their TTV-uncorrected counterparts were strong (mean r=0.94, range: 0.87-0.99) indicating that the ranking of cortical regions in each IDP change map was not substantially altered by exclusion or inclusion of TTV as a covariate when calculating IDP change (**Fig S4b**). In accordance with this phenomenon, the hierarchical structure of the correlation matrix between IDP change maps was also largely unaltered when excluding TTV as a covariate, with similarities amongst the IDP change maps being organized first by imaging modality and second by aneuploidy (**Fig S4c,d**). We also compared PCA outputs computed using the TTV corrected and uncorrected data and found a less striking similarity between XXY PC1 and XYY PC1 with regard to the PC scores; however, principal components other than the first one did not exhibit greater concordance (**Fig S6a,b**). Similarly, PC1 of XXY exhibited greater correlation with PC2 of XYY and T21 with regard to feature weights (**Fig S6c**). Despite these subtle differences, biological annotation of the average PC1 map using the Yeo-Krienen-17 parcellation identified a somatomotor subnetwork (MOT-1) and a ventral attention subnetwork (VAN-1) as showing enrichment for negative and positive PC1 scores, respectively — partly recapitulating what we had observed with the TTV-corrected PC1 results (**Fig S6d,f**).

## Discussion

Our study provides an unprecedentedly deep phenotypic analysis of neuroimaging changes in multiple GDDs by examining multimodal neuroimaging data in 3 different aneuploidy syndromes. This enriched dataset advances understanding of each individual aneuploidy considered; reveals organizing principles for regional cortical change across aneuploidies; and localizes cortical regions with shared vulnerability across aneuploidies. The cortical map of shared vulnerability to gene dosage change for three distinct supernumerary chromosomes — which is enriched for specific functional systems and cellulo-molecular signatures — significantly correlates with a map of shared cortical vulnerability to multiple behaviorally defined psychiatric disorders, thereby defining a key spatial axis of cortical organization for future translational study. We consider each output of our study in further detail below.

Most prior neuroimaging studies of XXY, XYY, and T21 have examined changes in GMV, CT, and SA, and we replicate previously reported findings. We identify folding changes as drivers of SA and GMV changes in both XXY and XYY and find evidence of a distinct morphometric signature in T21 where GMV changes showed mixed regional coherence with SA changes in posterior regions and CT changes in anterior regions. By extending beyond sMRI phenotypes, we add multiple new phenotypic layers to better understand cortical vulnerability in response to each of the aneuploidies studied. Salient examples include increased diffusivity in the temporal cortex and decreased diffusivity in the cingulate cortex for XXY, which may reflect altered myelin integrity. In addition, although we only find a statistically significant increase in anisotropy in the auditory association cortex for T21, this finding diverges from a previously reported decrease in anisotropy in a few cortical brain regions using the same T21 dataset^21^. For alterations across rsFMRI phenotypes, reduced ALFF in the inferior frontal cortex for XXY and XYY may reflect reduced activity or hypo-activation^22^. An important note is that the sparse statistically significant effects detected for DWI and rsFMRI were a result of the reduced power for these two modalities. Given that downstream analyses integrating multimodal data are contingent on continuous regional variation in effect size, it is imperative that future studies acquire larger multimodal sample sizes to increase power. Nonetheless, the diversity of regional changes observed across the different cortical phenotypes and aneuploidies implies highly variegated mechanisms for regional cortical change in each aneuploidy and shows how this complexity is missed by focusing on the most commonly measured sMRI phenotype alone.

By systematically comparing the spatial pattern of cortical change for all aneuploidy-phenotype combinations, our study pinpoints potential organizing principles for the patterning of cortical change in GDDs more broadly. For example, in the three aneuploidies considered here, we find that correlations are higher across spatial patterns of change across aneuploidies for a phenotype derived from a specific modality as opposed to within an aneuploidy, e.g. patterns of microstructural change in XXY correlate more strongly with patterns of microstructural change in XYY and T21 than with patterns of volumetric or local connectivity change in XXY. This organizing principle may have several explanations and implications. High correlations across phenotypes derived from the same modality are likely rooted in the phenotypes sharing a genetic architecture and capturing similar tissue properties^23^. This observed within-modality coherence helps to support phenotype selection in neuroimaging studies of additional GDDs by defining sets of phenotypes that tend to change together, enabling more efficient measurement of distinct patterns of change. This finding also suggests that there are dissociable mechanisms underpinning altered regional anatomy, white matter microstructure, and resting state function in humans, which is supported by past literature in mouse work and can shape downstream translational studies^24,25^.

The secondary organizing principle we detected — that regional change varies by aneuploidy type within each conjointly altered set of phenotypes — points to specificity of gene dosage change. Thus, even amongst aneuploidies that each alter the dosage of large gene sets with a theoretically greater potential for overlapping effects, we still observe differentiable spatial patterns of cortical change.

Lastly, we integrated information from our deep phenotypic analysis and revealed primary spatial axes of multimodal change for each aneuploidy. The motivation for compressing this information was to identify a latent dimension of change that may best summarize the spatial patterning of disruption identified across multiple phenotypes, and we extended this to identify regions that are maximally and minimally affected across all three aneuploidies. Regions that were convergently maximally affected included insular and ventromedial prefrontal areas that overlapped with ventral attention and somatomotor functional subnetworks. The observation of convergent aneuploidy effects in the insula is striking given that this same region shows overlapping anatomical changes across multiple BDDs^24^. Our finding of a clear spatial alignment between the average aneuploidy PC1 map and morphometric PC1 map in ENIGMA beckons the question of what biological features might underpin the apparently shared vulnerability of certain cortical regions to the diversity of risk factors captured by GDDs and BDDs. We tackle this question by evaluating alignment between our average aneuploidy PC1 map and spatial gene expression from the AHBA, which highlights the specific molecular signatures characterizing these shared regions of cortical change between different GDDs and BDDs. The enrichment of serotonergic pathways within these regions is particularly interesting given that the aneuploidies considered here increase risk for mental disorders that are associated with serotonergic dysfunction^27^, and recent work has also pointed to an association between the patterning of cross-disorder morphometric change in ENIGMA and the spatial distribution of the 5HT1-a and 5HT1-b serotonin receptors^16^. The specific enriched inhibitory interneuron cell class, In1, has been found to overlap with genes that are upregulated in cortical samples from individuals with ASD^28^, further supporting a potential mechanistic link between these aneuploidies and the BDDs they increase risk for. However, we note that these findings and interpretation are based only on the first principal component of cortical change in each aneuploidy, which captures only a fraction of all multimodal signals. Thus, there may be additional signatures of multimodal change that would align with other molecular and cellular processes.

Our findings must be considered in light of several caveats and limitations. First, although we include substantially more IDPs than previously studied in multiple GDDs, future work needs to expand even further on the set of phenotypes studied and include other GDDs. For example, more advanced DWI acquisitions would enable neurite-specific measures to be computed that are less susceptible to partial volume effects and noise, and novel methods are being developed to overcome differential regional signal loss in fMRI^29,30^. In addition, enhanced reproducibility and generalizability of results may require multiverse analytic approaches to be explored. For instance, the unthresholded FA maps for T21 in the current report differ from those generated in a previous study using a different processing pipeline^21^, suggesting that sub-threshold effects in small samples may be sensitive to differences in image processing pipelines and modeling choices. Second, with regard to the cross-aneuploidy comparisons and the potential generic mechanisms we identified, it is important to also consider aneuploidy-specific questions, such as which specific genes might be driving the dominant spatial profiles of multimodal change. Third, our cross-sectional study design does not allow us to capture how the cortical changes may unfold developmentally - both as it relates to individual aneuploidies or shared mechanisms. Relatedly, the AHBA analysis makes use of gene expression data in adults, so it may miss molecular and cellular mechanisms that are specific to atypical cortical organization in early development. Finally, it remains an open question as to whether variation in multimodal changes amongst carriers of the same GDD relate to variation in clinical severity, but it is unclear whether brain-behavior relationships are stronger in highly penetrant GDDs, and the rarity of the GDDs makes it difficult to amass an appropriately large sample size.

Notwithstanding the above limitations, our study secures a more comprehensive picture of cortical alterations across three distinct aneuploidies and emphasizes a shared backbone of cortical vulnerability amidst the diversity of effects for different IDPs in different GDDs. Strikingly, this shared signature of cortical vulnerability to aneuploidy aligns with the shared signature of cortical change amongst diverse behaviorally-defined disorders that occur at elevated rates in aneuploidies and other GDDs. The cortical gradient revealed by these shared effects represent a high-priority target for future translational research in basic and clinical neuroscience.

## Methods

### Participants

Our study includes a total of 341 participants aged 6-25 years from 3 case-control cohorts: XXY (total n = 191, 92 controls), XYY (total n=81, 47 controls), and T21 (total n=69, 41 controls). Participants were recruited through parent support organizations, the National Institute of Mental Health website, and the National Institutes of Health (NIH) Healthy Volunteer office. All 341 participants had sMRI data available, with subsets of each case-control cohort having DWI (total n=253; XXY case/control = 83/69; XXY case/control = 21/41; T21 case/control = 25/14) and rsFMRI data (total n=244; XXY case/control = 66/73; XYY case/control = 22/42; T21 case/control = 12/29). Participant characteristics are given in **Table 1** for the full sample, and are broken down by imaging modality in **Table S1**. All participants had normal radiological reports and no prior brain injuries. Complete information about the T21 sample can be found in ref^7^. All XXY and XYY participants were non mosaic with a genetic diagnosis confirmed by karyotype, and all controls were screened to exclude a history of neurodevelopmental or psychiatric disorders. For T21, all participants had clinically diagnosed T21, which was confirmed on clinical assessment and re-verified by karyotype in 20 participants who gave blood and by inspection of available karyotype reports in four of the remaining eight who were unable to give blood. This study was approved by the NIH Combined Neuroscience Institutional Review Board. All participants gave consent or assent, as appropriate, and all protocols were completed at the NIH Clinical Center in Bethesda, Maryland.

**Table 1.** Participant Characteristics for Full Sample. For each of the three modalities, we specify the sample size for cases and controls as well as the corresponding average age and standard deviation. We provide the FS IQ and SES for all subjects as well. T21 (trisomy 21) cases and controls included both males and females. Please see Table S1 for global measures across all IDPs and aneuploidies. FS IQ = Full Scale IQ; SES = Socioeconomic Status; SD = standard deviation.

|  |  | XXY |  | XYY |  | T21 |  |
| --- | --- | --- | --- | --- | --- | --- | --- |
|  |  | Case | Control | Case | Control | Case | Control |
| **Structural** | Participants | 99 | 92 | 34 | 47 | 28 | 41 |
|  | Age (SD) | 16.4 (4.8) | 16.2 (5.6) | 15.5 (5.3) | 14.0 (4.7) | 15.7 (5.5) | 16.1 (5.9) |
| **Diffusion** | Participants | 83 | 69 | 21 | 41 | 14 | 25 |
|  | Age (SD) | 16.3 (4.9) | 16.2 (5.4) | 16.1 (5.4) | 13.9 (4.8) | 17.6 (5.4) | 17.1 (6.2) |
| **Functional** | Participants | 66 | 73 | 22 | 42 | 12 | 29 |
|  | Age (SD) | 16.9 (4.2) | 17.9 (4.8) | 17.1 (4.5) | 14.7 (4.4) | 17.2 (4.7) | 18.5 (4.5) |
| **Demo** | FS IQ (SD) | 93.3 (12.2) | 115.1 (12.6) | 87.1 (12.4) | 118.5 (9.8) | 53.6 (14.6) | 116.3 (15.8) |
|  | SES (SD) | 47.3 (20.6) | 37.7 (17.4) | 54.9 (18.0) | 38.3 (15.4) | 41.0 (14.6) | 39.2 (21.5) |

### Neuroimaging Data Acquisition and Preprocessing

All sMRI (T1w-MP-RAGE), dWI (diffusion weighted imaging) and rsFMRI (10 minutes EPI for XXY and XYY; 5 minutes and 15 seconds EPI for T21) data were gathered using one MR750 3-Tesla scanner (General Electric, MA) for XXY and XYY and another GE 3-Tesla scanner for T21 with identical sequences within each case-control aneuploidy cohort (**Supplementary Materials**). For each cohort, T1w MRI and rs-fMRI scans were converted from DICOM to Nifti and organized according to the Brain Imaging Data Structure^31^ (BIDS) using heudiconv^32^. Each imaging modality was preprocessed as summarized below, with full methods in **Supplemental Materials**.

#### Structural MRI

We used the BIDS compatible Freesurfer pipeline version 7.1.0 to process each subject’s T1w MRI scan through the automated *recon-all* processing stream to generate the cortical surface reconstruction. This pipeline is extensively documented online (http://surfer.nmr.mgh.harvard.edu/)^33^. The cortical surface reconstruction steps were comprised of: skull stripping^34^; segmentation of cortical gray and white matter^35,36^; separation of the two hemispheres and subcortical structures^37–39^; and finally, construction of a smooth representation of the gray–white interface and the pial surface^40^. The following surface-based morphometric features were estimated at each of ~326k surface vertices using the FreeSurfer *mri_anatomical_stats* utility: CT, SA, GMV’ MeanCurv, GausCurv, CurvInd, and FoldInd. For each feature other than SA and GMV, vertex-level values were averaged to estimate regional values for all 360 cortical parcels in the multimodally-derived Human Connectome Project (HCP) regional parcellation^11^. For SA and GMV, the vertex-level values were summed to estimate regional values. For each cortical reconstruction we extracted the Euler number, a measure of topological complexity and image quality. Based on past guidelines all participants in this study had an sMRI-derived Euler number of greater than −217^41^, and individual-specific Euler numbers were included in all statistical models as a proxy of image quality (see below).

#### Diffusion weighted imaging

As previously described, DWI DICOM files were imported and converted to Nifti (using accompanying gradient files) through the ‘ImportDICOM’ function from TORTOISE^42,43^, which also extracted information about the b-vals and b-vecs. Subsequently, the following partially dissociable^44^ voxelwise diffusion tensor imaging (DTI) metrics were extracted using Tractoflow version 2.2.1^45^: FA, GA, MD, RD, AD, and TM (**Supplementary Materials**). Subject-specific maps were resampled to be in the space of the respective subject’s volumetric HCP parcellation in Freesurfer space (generated using the sMRI workflow), and voxel-wise estimates of the features were averaged within the corresponding region in the parcellation.

#### Resting-state functional MRI

All rs-fMRI scans were processed using fMRIPrep version 20.2.0., which is based on *Nipype* version 1.5.1^46–48^. Thorough descriptions of fMRIPrep’s processing steps are provided by the creators of the software under a CC0 license and are reproduced in Supplementary Materials. As per [Esteban et al, 2019], fMRIPrep accepts one T1w image and one BOLD series per subject and generates a preprocessed BOLD run in MNI 152 template space, along with a set of confound estimates for nuisance regression. The pre-processed BOLD scan and relevant confounds were subsequently submitted to the extensible Connectivity Pipeline (XCP) Engine to denoise the fMRI signal using anatomical component-based correction^49^ (**Supplementary Materials**). XCPEngine was used to compute two voxel-wise metrics of local connectivity - ALFF and ReHo - for participants with average head motion <0.3 mm/TR. The ALFF calculation consisted of computing a power spectrum at each voxel, then integrating over the 0.01-0.08 Hz passband of the power spectrum. Regional homogeneity for each voxel was computed as Kendall’s W (coefficient of concordance) among the timeseries of the voxel and its 26 neighboring voxels. Using the volumetric HCP parcellation, XCPEngine computes the mean value of each voxelwise metric across all ROIs of the parcellation. rsFMRI data are subject to well-described signal losses in certain areas of the brain^50,51^, and we removed 16/360 HCP ROIs from all subsequent analyses based on their missing rs-fMRI data for 20 or more subjects (**FigS9**).

### Computing regional IDP changes in each aneuploidy

The aforementioned preprocessing steps resulted in complete person-specific measures of 15 IDPs for 344 cortical ROIs in each of the three case-control cohorts (one for each GDD). In each of the cohorts, available person-specific measures were used to compute 15 aneuploidy-specific ΔIDP maps through the workflow depicted in **Figure 1**. For each IDP and cohort, we first z-scored each ROI’s IDP values across individuals and then modeled the effect of group status (case versus control) on the IDP while controlling for age and total tissue volume (TTV). Total tissue volume was included as a covariate in main analyses given the known robust size reductions in XXY and T21 combined with mean TTV increases in XYY^1,7^ to protect against the ROI-level GDD comparisons being trivially colored by global effects [especially for ROI sMRI measures that scale closely with TTV^52^]. However, all analyses were repeated without TTV covariation, as detailed in **Supplementary Materials** and referred to in **Results**. Additional covariates were the Euler number as well as framewise displacement for all rsFMRI-derived IDPs and sex for all IDPs in the T21 case-control cohort. Case-control differences were estimated as the standardized beta coefficients (henceforth “effect sizes”) for the group covariate. To identify statistically-significant regional IDP changes, FDR correction was performed using the Benjamini-Hochberg approach for each set of 344 p-values relating to a specific IDP change in a specific aneuploidy (i.e., treating each GDD-I DP combination as a separate family of tests).

### Defining organizing-principles of cortical change across IDPs, regions and aneuploidies

We estimated the pairwise similarity between all 45 un-thresholded ΔIDP maps (15 IDPs for each of 3 aneuploidies) based on their spatial correlation across all 344 ROIs. The resulting symmetrical 45*45 correlation matrix was characterized using two complementary approaches. First, we used hierarchical clustering to describe the broad structure of this correlation matrix, paying particular attention to whether ΔIDP groups were organized primarily by modality or aneuploidy type. Second, we used a previously reported spatial permutation framework to identify statistically significant inter-map similarities amongst all 990 unique pairwise comparisons while accounting for spatial autocorrelations within maps and correcting for multiple comparisons (**Supplementary Methods**)^12,53^.

### Defining and characterizing principal components of multimodal cortical change in each aneuploidy

We next used principal component analysis (PCA) to provide an efficient summary representation of multimodal change for each aneuploidy, defined by ROI scores for the first principal component (PC1) of the 344*15 ΔIDP matrix and associated feature loadings for this PC. We compared PC1 across aneuploidies based on (i) the cross-ROI spatial correlation in PC1 scores, and (ii) the cross-IDP correlation in feature loadings. To aid cross-aneuploidy comparisons, the polarity of these first PCs were set to maximize positive inter-aneuploidy PC1 map correlation. We focused our main analyses on PC1, but maps of ROI scores and details of IDP loadings are provided for the first five PCs in each aneuploidy (explaining a cumulative variance >80%) as **Supplementary Materials**.

To put PC1 in broader biological context, we compared the regional distribution of PC1 scores in each aneuploidy against canonical parcellations of the human cortex based on resting state functional connectivity networks^14^ (Yeo-Krienen-17 atlas, 17 networks). For each aneuploidy, we first asked if the observed omnibus effect of each cortical parcellation on PC1 scores exceeded the distribution of effects from 1000 spatial permutations of the PC1 map (**Supplementary Materials**). Where this test indicated a significant alignment between PC1 and the parcellation, we conducted post hoc tests comparing observed PC1 scores for each component of the parcellation against their respective null distributions to identify specific functional networks that were significantly enriched (at p<0.05) for extreme PC1 scores of multimodal cortical change.

### Assessing multimodal convergence across aneuploidies

The PC1 maps of multimodal change in each aneuploidy (see above) were used to test for potential regions of convergent cortical vulnerability or resilience across aneuploidies. To assess convergent vulnerability, we defined the upper and lower decile of ROIs by PC1 score within each aneuploidy and generated a single cross-aneuploidy conjunction map for each of these extremes to show regions included in 1, 2 or all 3 groups. For each of the two PC1 extremes, we considered regions impacted by 2 or more aneuploidies to show convergent vulnerability. Similarly, regions of convergent resilience were defined as those that fell in the middle decile of PC1 scores for 2 or more aneuploidies. The three groups of convergence were subsequently annotated using the same parcellations applied to annotate each aneuploidy-specific PC1^14^.

### Assessing spatial alignment with the principal component of morphometric change in behaviorally defined psychiatric disorders

The ENIGMA toolbox^54^ contains case-control maps for cortical thickness and surface area data across 68 brain regions defined using the Desikan-Killiany (DK) parcellation (**Supplementary Materials**). We specifically obtained a total of 21 case-control maps spanning autism spectrum disorder (ASD), attention deficit hyperactivity disorder (ADHD), bipolar disorder (BD), schizophrenia (SCZ), major depressive disorder (MDD), and obsessive compulsive disorder (OCD), and we applied PCA to identify the first principal component (ENIGMA-PC1) of morphometric change across the 21 maps. We sought to compare the spatial alignment between the ENIGMA-PC1 map with the average PC1 map across XXY, XYY, and T21 (Aneuploidy-PC1). We thus mapped the HCP ROIs onto the Freesurfer 5 cortical surface vertices, and averaged the PC1 vertex-wise scores across the DK ROIs while ignoring surface vertices corresponding to the HCP ROIs excluded from our analysis. We then computed the Pearson correlation between the ENIGMA-PC1 and Pearson-PC1 maps and performed a spin test to assess significance.

### Gene-category enrichment analysis of the cross-disorder aneuploidy map

We used the Allen Human Brain Atlas to compute gene expression for 15,633 genes across the HCP-Glasser brain regions (**Supplementary Materials**) for performing gene category enrichment analysis (GCEA) of the Aneuploidy-PC1 map. To overcome recently reported non-negligible false positive rates when applying GCEA, we used ABAnnotate^19^, a toolbox that applies a nonparametric method introduced in ref^18^ to estimate significant correlations relative to an ensemble of randomized phenotypes. In keeping with the previous analyses, we generated the randomized phenotypes by spinning the empirical Aneuploidy-PC1 map 1000 times. ABAnnotate facilitates GCEA using various gene-category datasets, and we used two of the included datasets: “GO-biologicalProcessProp-discrete” for gene-biological process annotations^55–58^ and “PsychEncode-cellTypesTPM-discrete” for neuronal cell type marker genes^59,60^. ABAnnotate looks for genes with a positive correlation with a given phenotype, so we accounted for the polarity of the PC1 map by applying the toolbox both to the original Aneuploidy-PC1 map and the flipped Aneuploidy-PC1 map. Separately for each of the three GCEA analyses, the toolbox first computes the enrichment score for each category by averaging across the Spearman rank correlation between each gene in a category and the Aneuploidy-PC1 map. A null distribution of category enrichment scores is computed relative to the null Aneuploidy-PC1 maps, and a p-value is estimated by comparing the empirical category score to the null distribution and then false-discovery rate corrected (FDR < 0.05).

## Code availability

All analyses have been made available at https://github.com/llevitis/NIH_Multimodal_SCA_Phenotypes.

## Acknowledgments

The authors would like to thank all the participants and their families for their generous involvement in this study. This study was supported by the intramural research program of the National Institute of Mental Health (project funding: 1ZIAMH002949-03; clinical trials identifier: NCT00001246; clinicaltrials.gov; protocol: 89-M-0006) and the National Institute of Child Health and Disease (R01HD100298). The authors acknowledge members of the UCL POND group (http://pond.cs.ucl.ac.uk) for feedback and input received during group discussions.

## Author contributions

Conceptualization, writing - original draft, EL, AR, DA; project administration, EL, AR, DA; methodology, software, and formal analysis, EL, SL, AR, DA; visualization, EL; data collection, LC, ET, AW, EW, JB, JS, NL, FL; funding acquisition, AR; writing - review & editing, all authors; supervision, AR, DA.

## Resource Availability

Further information and requests for resources should be directed to Dr. Armin Raznahan.

**Supplemental materials**

## Legends for any Supplemental Excel Tables or files

***Figure S1. Quality control workflow**. The sMRI workflow is in yellow, rs-FMRI in purple, and DWI in green. T1w scans were first assessed by a radiologist for any abnormalities, they were then processed using MRIQC to identify any scans with outlying image quality metrics, and the Freesurfer-generated Euler number was used as a final QC step.*

***Figure S2. IDP maps for both hemispheres with TTV correction.** a) Unthresholded effect sizes where blue colors denote a decrease in cases relative to controls and red colors denote an increase in cases relative to controls. b) Thresholded effect sizes where all regions of interest colored in red exhibit statistically significant increases in cases relative to controls, and all regions of interest colored in blue exhibit statistically significant decreases in cases relative to controls.*

***Figure S3. IDP maps for both hemispheres without TTV correction.** a) Unthresholded effect sizes where blue colors denote a decrease in cases relative to controls and red colors denote an increase in cases relative to controls. b) Thresholded effect sizes where all regions of interest colored in red exhibit statistically significant increases in cases relative to controls, and all regions of interest colored in blue exhibit statistically significant decreases in cases relative to controls.*

***Figure S4. Visualizing aneuploidy specific ROI by ΔIDP maps and cross-phenotypic relationships without correction for total tissue volume.** a) ROI by ΔIDP matrices for each phenotype, where each cell represents the standardized beta coefficient for group effect on a particular aneuploidy at a given brain region. b) Correlation of ROI by ΔIDP maps across all aneuploidies with and without correction for total tissue volume. c) Hierarchical clustering matrix of the concatenated 344 by 45 ROI by ΔIDP matrix. Each row and column corresponds to an aneuploidy-IDP combination. d) Hierarchical clustering matrix of the difference between the cross-correlation matrix generated using the ROI by ΔIDP matrices with TTV correction minus the ROI by ΔIDP matrices without TTV correction.*

***Figure S5. PCA results for the analytical stream with TTV correction.** a) Scores for the first five principal components projected onto the surface for each aneuploidy. b) Feature weights for the first five principal components. c) Cross-correlation of each aneuploidy-PC score combination. d) Cross-correlation of each aneuploidy-PC feature combination.*

***Fig S6. PCA results for the analytical stream without TTV correction. (a)** Scores for the first five principal components projected onto the surface for each aneuploidy. **(b)** Cross-correlation of each aneuploidy-PC score combination both with and without TTV correction. **(c)** Cross-correlation of each aneuploidy-PC feature combination both with and without TTV correction. **(d)** Average PC1 map (Aneuploidy-PC1) projected onto the cortical surface (in the HCP/Glasser atlas parcellation). (**e**) Conjunction map showing ROIs which sit at the top (red), middle (greens) and bottom (blues) deciles of PC1 scores across aneuploidy/GDD. (**f**) Enrichment of functional connectivity modules defined by Yeo-Krienen-17 (top cortical surfaces, mapped to the Glasser parcellation), for extreme PC1 scores within the Aneuploidy-PC1 map.*

***Fig S7. Empirical versus null F-statistics across aneuploidy PC1 maps and annotations.** a) Results using data computed with TTV correction. b) Results using data computed without TTV correction. For both panels, the null distributions of the F-statistics are colored in red, and the empirical F-statistic is in black.*

***Fig S8. Enrichment of functional connectivity modules defined by Yeo-Krienen-17 for extreme PC1 scores within XXY, XYY, and T21 PC1 maps.** a) Cortical surfaces, mapped to the Glasser parcellation. b) Results using PC1 outputs generated using TTV corrected data. c) Results using PC1 outputs generated using TTV uncorrected data.*

***Fig S9. rs-FMRI signal loss.** Using the XXY rsfMRI data, we show the number of participants who exhibit signal loss for particular regions of interest following processing with xcpEngine. ROIs lacking coverage for at least 20 participants were bilaterally excluded from the analysis for each aneuploidy.*

***Table S1. Global measures for all IDPs, TTV, FD, and Euler number.** For area and volume we display the total value for each measure across the 344 included HCP ROIs averaged across subjects whereas for the remaining phenotypes, we display the average value for each measure across the 344 included HCP ROIs averaged across subjects. FD is computed using subjects who have functional MRI data while TTV and euler_mean_bh are for all subjects. Euler_mean_bh = Euler number; TTV = total tissue volume; FD = framewise displacement.*

***Table S2. Summary table for effect sizes for all aneuploidies with and without TTV correction.** Each effect size has an accompanying FDR corrected q-value (see attached document: supp_xxy_xyy_t21_effect_sizes.xlsx).*

***Table S3. Summary table for PCA results for all aneuploidies with and without TTV correction.** The sheets labeled “pca_scores_withTTV” and “pca_features_withTTV” contain the principal component scores and feature weights across the top five PCs for each aneuploidy following the processing pipeline with TTV correction. The sheets labeled “pca_scores_withoutTTV” and “pca_features_withoutTTV” contain the principal component scores and feature weights for each aneuploidy following the processing pipeline without TTV correction.*

***Table S4. Summary table for GCEA results generated using ABAnnotate.** The sheets “aneuploidyPC1_BP” and “aneuploidyPC1reverse_BP” contain the GCEA results for the gene ontology (GO) biological processes and the original and flipped PC1 maps, respectively. The sheets “aneuploidyPC1_celltypes” and “aneuploidyPC1reverse_celltypes” contain the GCEA results for the cell types and the original and flipped PC1 maps, respectively.*

## References

1. Raznahan A, Lee NR, Greenstein D, et al. Globally Divergent but Locally Convergent X- and Y-Chromosome Influences on Cortical Development. Cereb Cortex. 2016;26(1):70–79. doi:10.1093/cercor/bhu174

2. Moreau CA, Urchs SGW, Kuldeep K, et al. Mutations associated with neuropsychiatric conditions delineate functional brain connectivity dimensions contributing to autism and schizophrenia. Nat Commun. 2020;11(1):5272. doi:10.1038/s41467-020-18997-2

3. Seidlitz J, Nadig A, Liu S, et al. Transcriptomic and cellular decoding of regional brain vulnerability to neurogenetic disorders. Nat Commun. 2020;11(1):3358. doi:10.1038/s41467-020-17051-5

4. Modenato C, Kumar K, Moreau C, et al. Effects of eight neuropsychiatric copy number variants on human brain structure. Transl Psychiatry. 2021;11(1):399. doi:10.1038/s41398-021-01490-9

5. Rau S, Whitman ET, Schauder K, et al. Patterns of Psychopathology and Cognition in Sex Chromosome Aneuploidy. In Review; 2021. doi:10.21203/rs.3.rs-543874/v1

6. Lepage JF, Hong DS, Raman M, et al. Brain morphology in children with 47,XYY syndrome: a voxel- and surface-based morphometric study: Brain morphology in XYY. Genes Brain Behav. 2014;13(2):127–134. doi:10.1111/gbb.12107

7. Lee NR, Adeyemi EI, Lin A, et al. Dissociations in Cortical Morphometry in Youth with Down Syndrome: Evidence for Reduced Surface Area but Increased Thickness. Cereb Cortex. 2016;26(7):2982–2990. doi:10.1093/cercor/bhv107

8. Hamner T, Udhnani MD, Osipowicz KZ, Lee NR. Pediatric Brain Development in Down Syndrome: A Field in Its Infancy. J Int Neuropsychol Soc. 2018;24(9):966–976. doi:10.1017/S1355617718000206

9. Nielsen J, Wohlert M. Chromosome abnormalities found among 34910 newborn children: results from a 13-year incidence study in ◆rhus, Denmark. Hum Genet. 1991;87(1):81–83. doi:10.1007/BF01213097

10. Vicari S. Motor Development and Neuropsychological Patterns in Persons with Down Syndrome. Behav Genet. 2006;36(3):355–364. doi:10.1007/s10519-006-9057-8

11. Glasser MF, Coalson TS, Robinson EC, et al. A multi-modal parcellation of human cerebral cortex. Nature. 2016;536(7615):171–178. doi:10.1038/nature18933

12. Alexander-Bloch A, Giedd JN, Bullmore E. Imaging structural co-variance between human brain regions. Nat Rev Neurosci. 2013;14(5):322–336. doi:10.1038/nrn3465

13. Váša F, Seidlitz J, Romero-Garcia R, et al. Adolescent Tuning of Association Cortex in Human Structural Brain Networks. Cereb Cortex. 2018;28(1):281–294. doi:10.1093/cercor/bhx249

14. Thomas Yeo BT, Krienen FM, Sepulcre J, et al. The organization of the human cerebral cortex estimated by intrinsic functional connectivity. J Neurophysiol. 2011;106(3):1125–1165. doi:10.1152/jn.00338.2011

15. Opel N, Goltermann J, Hermesdorf M, Berger K, Baune BT, Dannlowski U. Cross-Disorder Analysis of Brain Structural Abnormalities in Six Major Psychiatric Disorders: A Secondary Analysis of Mega- and Meta-analytical Findings From the ENIGMA Consortium. Biol Psychiatry. 2020;88(9):678–686. doi:10.1016/j.biopsych.2020.04.027

16. Park B yong, Kebets V, Larivière S, et al. Multiscale neural gradients reflect transdiagnostic effects of major psychiatric conditions on cortical morphology. Commun Biol. 2022;5(1):1024. doi:10.1038/s42003-022-03963-z

17. Hawrylycz M, Miller JA, Menon V, et al. Canonical genetic signatures of the adult human brain. Nat Neurosci. 2015;18(12):1832–1844. doi:10.1038/nn.4171

18. Fulcher BD, Arnatkeviciute A, Fornito A. Overcoming false-positive gene-category enrichment in the analysis of spatially resolved transcriptomic brain atlas data. Nat Commun. 2021;12(1):2669. doi:10.1038/s41467-021-22862-1

19. Lotter LD, Dukart J, Fulcher BD. ABAnnotate: A toolbox for ensemble-based multimodal gene-category enrichment analysis of human neuroimaging data. Published online April 15, 2022. doi:10.5281/ZENODO.6463329

20. Reijnders MJMF, Waterhouse RM. Summary Visualizations of Gene Ontology Terms With GO-Figure! Front Bioinforma. 2021;1:638255. doi:10.3389/fbinf.2021.638255

21. Lee NR, Nayak A, Irfanoglu MO, et al. Hypoplasia of cerebellar afferent networks in Down syndrome revealed by DTI-driven tensor based morphometry. Sci Rep. 2020;10(1):5447. doi:10.1038/s41598-020-61799-1

22. Aiello M, Salvatore E, Cachia A, et al. Relationship between simultaneously acquired resting-state regional cerebral glucose metabolism and functional MRI: a PET/MR hybrid scanner study. NeuroImage. 2015;113:111–121. doi:10.1016/j.neuroimage.2015.03.017

23. van der Meer D, Frei O, Kaufmann T, et al. Understanding the genetic determinants of the brain with MOSTest. Nat Commun. 2020;11(1):3512. doi:10.1038/s41467-020-17368-1

24. Ellegood J, Anagnostou E, Babineau BA, et al. Clustering autism: using neuroanatomical differences in 26 mouse models to gain insight into the heterogeneity. Mol Psychiatry. 2015;20(1):118–125. doi:10.1038/mp.2014.98

25. Zerbi V, Pagani M, Markicevic M, et al. Brain mapping across 16 autism mouse models reveals a spectrum of functional connectivity subtypes. Mol Psychiatry. 2021;26(12):7610–7620. doi:10.1038/s41380-021-01245-4

26. Goodkind M, Eickhoff SB, Oathes DJ, et al. Identification of a Common Neurobiological Substrate for Mental Illness. JAMA Psychiatry. 2015;72(4):305. doi:10.1001/jamapsychiatry.2014.2206

27. Marazziti D. Understanding the role of serotonin in psychiatric diseases. F1000Research. 2017;6:180. doi:10.12688/f1000research.10094.1

28. Wang P, Zhao D, Lachman HM, Zheng D. Enriched expression of genes associated with autism spectrum disorders in human inhibitory neurons. Transl Psychiatry. 2018;8(1):13. doi:10.1038/s41398-017-0058-6

29. Zhang H, Schneider T, Wheeler-Kingshott CA, Alexander DC. NODDI: Practical in vivo neurite orientation dispersion and density imaging of the human brain. NeuroImage. 2012;61(4):1000–1016. doi:10.1016/j.neuroimage.2012.03.072

30. Yan Y, Dahmani L, Ren J, et al. Reconstructing lost BOLD signal in individual participants using deep machine learning. Nat Commun. 2020;11(1):5046. doi:10.1038/s41467-020-18823-9

31. Gorgolewski KJ, Auer T, Calhoun VD, et al. The brain imaging data structure, a format for organizing and describing outputs of neuroimaging experiments. Sci Data. 2016;3(1):160044. doi:10.1038/sdata.2016.44

32. Halchenko Y, Goncalves M, Castello MVDO, et al. nipy/heudiconv v0.9.0. Published online December 23, 2020. doi:10.5281/ZENODO.4390433

33. Fischl B. FreeSurfer. NeuroImage. 2012;62(2):774–781. doi:10.1016/j.neuroimage.2012.01.021

34. Ségonne F, Dale AM, Busa E, et al. A hybrid approach to the skull stripping problem in MRI. NeuroImage. 2004;22(3):1060–1075. doi:10.1016/j.neuroimage.2004.03.032

35. Fischl B, Liu A, Dale AM. Automated manifold surgery: constructing geometrically accurate and topologically correct models of the human cerebral cortex. IEEE Trans Med Imaging. 2001;20(1):70–80. doi:10.1109/42.906426

36. Segonne F, Pacheco J, Fischl B. Geometrically Accurate Topology-Correction of Cortical Surfaces Using Nonseparating Loops. IEEE Trans Med Imaging. 2007;26(4):518–529. doi:10.1109/TMI.2006.887364

37. Fischl B, Salat DH, Busa E, et al. Whole Brain Segmentation. Neuron. 2002;33(3):341–355. doi:10.1016/S0896-6273(02)00569-X

38. Fischl B. Automatically Parcellating the Human Cerebral Cortex. Cereb Cortex. 2004;14(1):11–22. doi:10.1093/cercor/bhg087

39. Dale AM, Fischl B, Sereno MI. Cortical Surface-Based Analysis. NeuroImage. 1999;9(2):179–194. doi:10.1006/nimg.1998.0395

40. Fischl B, Sereno MI, Dale AM. Cortical Surface-Based Analysis. NeuroImage. 1999;9(2):195–207. doi:10.1006/nimg.1998.0396

41. Rosen AFG, Roalf DR, Ruparel K, et al. Quantitative assessment of structural image quality. Neuroimage. 2018;169:407–418. doi:10.1016/j.neuroimage.2017.12.059

42. Pierpaoli C. TORTOISE: an integrated software package for processing of diffusion MRI data. Conference presented at: ISMRM 18th Annual Meeting; 2010; Stockholm, Sweden.

43. Irfanoglu MO. TORTOISEv3: Improvements and New Features of the NIH Diffusion MRI Processing Pipeline,. Abstract presented at: ISMRM 25th annual meeting; Honolulu, Hawaii.

44. Chamberland M, Raven EP, Genc S, et al. Dimensionality reduction of diffusion MRI measures for improved tractometry of the human brain. Neuroimage. 2019;200:89–100. doi:10.1016/j.neuroimage.2019.06.020

45. Theaud G, Houde JC, Boré A, Rheault F, Morency F, Descoteaux M. TractoFlow: A robust, efficient and reproducible diffusion MRI pipeline leveraging Nextflow & Singularity. Neuroimage. 2020;218:116889. doi:10.1016/j.neuroimage.2020.116889

46. Gorgolewski K, Burns CD, Madison C, et al. Nipype: A Flexible, Lightweight and Extensible Neuroimaging Data Processing Framework in Python. Front Neuroinformatics. 2011;5. doi:10.3389/fninf.2011.00013

47. Esteban, Oscar, Markiewicz, Christopher J., Burns, Christopher, et al. nipy/nipype: 1.7.0. Published online October 20, 2021. doi:10.5281/ZENODO.596855

48. Esteban O, Markiewicz CJ, Blair RW, et al. fMRIPrep: a robust preprocessing pipeline for functional MRI. Nat Methods. 2019;16(1):111–116. doi:10.1038/s41592-018-0235-4

49. Ciric R, Rosen AFG, Erus G, et al. Mitigating head motion artifact in functional connectivity MRI. Nat Protoc. 2018;13(12):2801–2826. doi:10.1038/s41596-018-0065-y

50. Jezzard P, Balaban RS. Correction for geometric distortion in echo planar images from B0 field variations. Magn Reson Med. 1995;34(1):65–73. doi:10.1002/mrm.1910340111

51. Norris DG, Zysset S, Mildner T, Wiggins CJ. An investigation of the value of spin-echo-based fMRI using a Stroop color-word matching task and EPI at 3 T. NeuroImage. 2002;15(3):719–726. doi:10.1006/nimg.2001.1005

52. Williams CM, Peyre H, Toro R, Ramus F. Neuroanatomical norms in the UK Biobank: The impact of allometric scaling, sex, and age. Hum Brain Mapp. 2021;42(14):4623–4642. doi:10.1002/hbm.25572

53. Váša F, Seidlitz J, Romero-Garcia R, et al. Adolescent Tuning of Association Cortex in Human Structural Brain Networks. Cereb Cortex. 2018;28(1):281–294. doi:10.1093/cercor/bhx249

54. Larivière S, Paquola C, Park B yong, et al. The ENIGMA Toolbox: multiscale neural contextualization of multisite neuroimaging datasets. Nat Methods. 2021;18(7):698–700. doi:10.1038/s41592-021-01186-4

55. Ashburner M, Ball CA, Blake JA, et al. Gene Ontology: tool for the unification of biology. Nat Genet. 2000;25(1):25–29. doi:10.1038/75556

56. The Gene Ontology Consortium, Carbon S, Douglass E, et al. The Gene Ontology resource: enriching a GOld mine. Nucleic Acids Res. 2021;49(D1):D325–D334. doi:10.1093/nar/gkaa1113

57. Huang DW, Sherman BT, Lempicki RA. Systematic and integrative analysis of large gene lists using DAVID bioinformatics resources. Nat Protoc. 2009;4(1):44–57. doi:10.1038/nprot.2008.211

58. Huang DW, Sherman BT, Lempicki RA. Bioinformatics enrichment tools: paths toward the comprehensive functional analysis of large gene lists. Nucleic Acids Res. 2009;37(1):1–13. doi:10.1093/nar/gkn923

59. Darmanis S, Sloan SA, Zhang Y, et al. A survey of human brain transcriptome diversity at the single cell level. Proc Natl Acad Sci. 2015;112(23):7285–7290. doi:10.1073/pnas.1507125112

60. Lake BB, Ai R, Kaeser GE, et al. Neuronal subtypes and diversity revealed by single-nucleus RNA sequencing of the human brain. Science. 2016;352(6293):1586–1590. doi:10.1126/science.aaf1204

