## Supplemental Materials for "The variegation of human brain vulnerability to rare genetic disorders and convergence with behaviorally defined disorders"

### Supplementary Materials

#### 1. Methods

##### 1.1 Neuroimaging data acquisition

High-resolution MP-RAGE T1-weighted structural magnetic resonance images (sMRI) were collected using the same protocol for participants in XXY and XYY case-control cohorts (172 contiguous sagittal slices with 256 x 256 in-plane matrix and 1 mm slice thickness yielding 1 mm isotropic voxels). For the T21 case-control cohort, the T1w images were collected with a 3D gradient echo sequence with the following protocol: 224 x 224 acquisition in-plane matrix, resolution of 0.94 x 0.94 x 1.2 mm, 128 slices, flip angle = 12°; field of view [FOV] = 240 mm.

Spontaneous slowly fluctuating brain activity was measured during rsFMRI through an echo-planar imaging (EPI) sequence while participants lay still and focused on a fixation cross. For the XXY and XYY case-control cohorts, the sequence was 10 min (repetition time of 2 s, echo time=0.03 s, flip angle=60°, 41 interleaved axial slices per volume, 3 mm slice thickness in a 216 mm x 216 mm acquisition matrix, single voxel=3 mm isotropic). For T21 case-control cohort, the sequence was 5 min (repetition time of 2.5 s, echo time=0.027 s, flip angle=90°, 44 interleaved axial slices per volume, 2.8 mm slice thickness in a 64 mm x 64 mm acquisition matrix, single voxel=2.8 mm).

The diffusion weighted imaging (DWI) acquisition differed across all the cohorts. The XXY case-control data was acquired using a reverse phase-encoding protocol (AP and PA) consisting of 1 unweighted B0 image, 3 gradient directions with a b-value = 200 mm/s2, 3 with a b-value = 500 mm/s2, and 38 with a b-value = 1000 mm/s2. This protocol also had these attributes: echo time = 0.085 s, repetition time = 9.947 s, 128 mm x 128 mm acquisition matrix, 2 mm slice thickness. The XYY and T21 case-control data were acquired with only AP phase-encoded DWI images. The XYY data consisted of 80 volumes with 10 b = 0, 10 b = 300, and 60 b = 1100 mm/s^2. The T21 data consisted of 60 volumes with 6 b=0, 12 b = 300, and 42 b = 1100 mm/s^2. The resolution was 2.5 mm isotropic zero-filled at the scanner to 1.87 × 1.87 × 2.5 mm.

##### 1.2 Diffusion MRI processing

The required inputs included: a DWI scan, b-values, b-vectors, an optional reversed phase encoded b0 (rev_b0.nii.gz) and T1-weighted image (t1.nii.gz) . For the XXY case-control dataset, we also supplied the reversed phase encoded b0 image to correct distortion due to diffusion acquisition. For XYY and T21 case-control datasets, we were not able to correct for susceptibility-induced distortion due to the reversed phase encoded b0 image not having been acquired. Tractoflow was used to perform all major processing steps, including preprocessing of structural and diffusion weighted images as well as the computation of DTI metric maps. Like the aforementioned BIDS-compatible pipelines, this toolbox utilizes functions from previously published neuroimaging software (e.g., FSL, MRtrix3, ANTs, and DIPY). Please see the original publication for an in depth description of all processing steps.

**1.3 Resting-state functional MRI processing**

Thorough descriptions of fMRIPrep's processing steps are provided by the creators of the software under a CC0 license and are reproduced here. Each T1w scan was corrected for intensity non-uniformity (INU) with N4BiasFieldCorrection[1], distributed with ANTs 2.3.3[2]. The T1w-reference was then skull-stripped with a *Nipype* implementation of the antsBrainExtraction.sh workflow (from ANTs), using OASIS30ANTs as target template. Brain tissue segmentation of cerebrospinal fluid (CSF), white-matter (WM) and gray-matter (GM) was performed on the brain-extracted T1w using fast (FSL 5.0.9)[3]. A T1w-reference map was computed after registration of T1w images (after INU-correction) using mri_robust_template, and brain surfaces were reconstructed using recon-all (FreeSurfer 6.0.1)[4]. The brain mask estimated previously was refined with a custom variation of the method to reconcile ANTs-derived and FreeSurfer-derived segmentations of the cortical gray-matter of Mindboggle[5]. Volume-based spatial normalization to one standard space (MNI152NLin2009cAsym) was performed through nonlinear registration with antsRegistration (ANTs 2.3.3), using brain-extracted versions of both T1w reference and the T1w template. The following template was selected for spatial normalization: *ICBM 152 Nonlinear Asymmetrical template version 2009c*[6].

For each of the BOLD runs, the following preprocessing was performed. First, a reference volume and its skull-stripped version were generated using a custom methodology of *fMRIPrep*. A deformation field to correct for susceptibility distortions was estimated based on *fMRIPrep*'s *fieldmap-less* approach. The deformation field is that resulting from co-registering the BOLD reference to the same-subject T1w-reference with its intensity inverted[7,8]. Registration is performed with antsRegistration, and based on the estimated susceptibility distortion, a corrected EPI (echo-planar imaging) reference was calculated for a more accurate co-registration with the anatomical reference. The BOLD reference was then co-registered to the T1w reference using bbregister (FreeSurfer) which implements boundary-based registration, and six degrees of freedom were used (Greve and Fischl 2009). Head-motion parameters with respect to the BOLD reference (transformation matrices, and six corresponding rotation and translation parameters) were estimated before any spatiotemporal filtering using FSL MCFLIRT. The BOLD time-series (including slice-timing correction when applied) were resampled onto their original, native space by applying a single, composite transform to correct for head-motion and susceptibility distortions. This BOLD time-series was then resampled into standard *MNI152NLin2009cAsym space* and used for calculating confounding time-series.

The pre-processed BOLD scan and relevant confounds were subsequently submitted to the eXtensible Connectivity Pipeline (XCP) Engine to denoise the fMRI signal and estimate measures of local functional connectivity[9]. Specifically, the anatomical component-based correction (aCompCor) processing stream was used to denoise the images[10]. aCompCor identifies sources of signal variance in white matter and cerebrospinal fluid using principal component analysis, and the principal components explaining 50% of the variance in the aforementioned signal are included in the denoising model, along with motion estimates and their derivatives.

##### 1.4 Quality control

Quality control (QC) was carried out separately for each imaging modality, and we report participant numbers at each modality-specific QC stage in **Fig S1**.

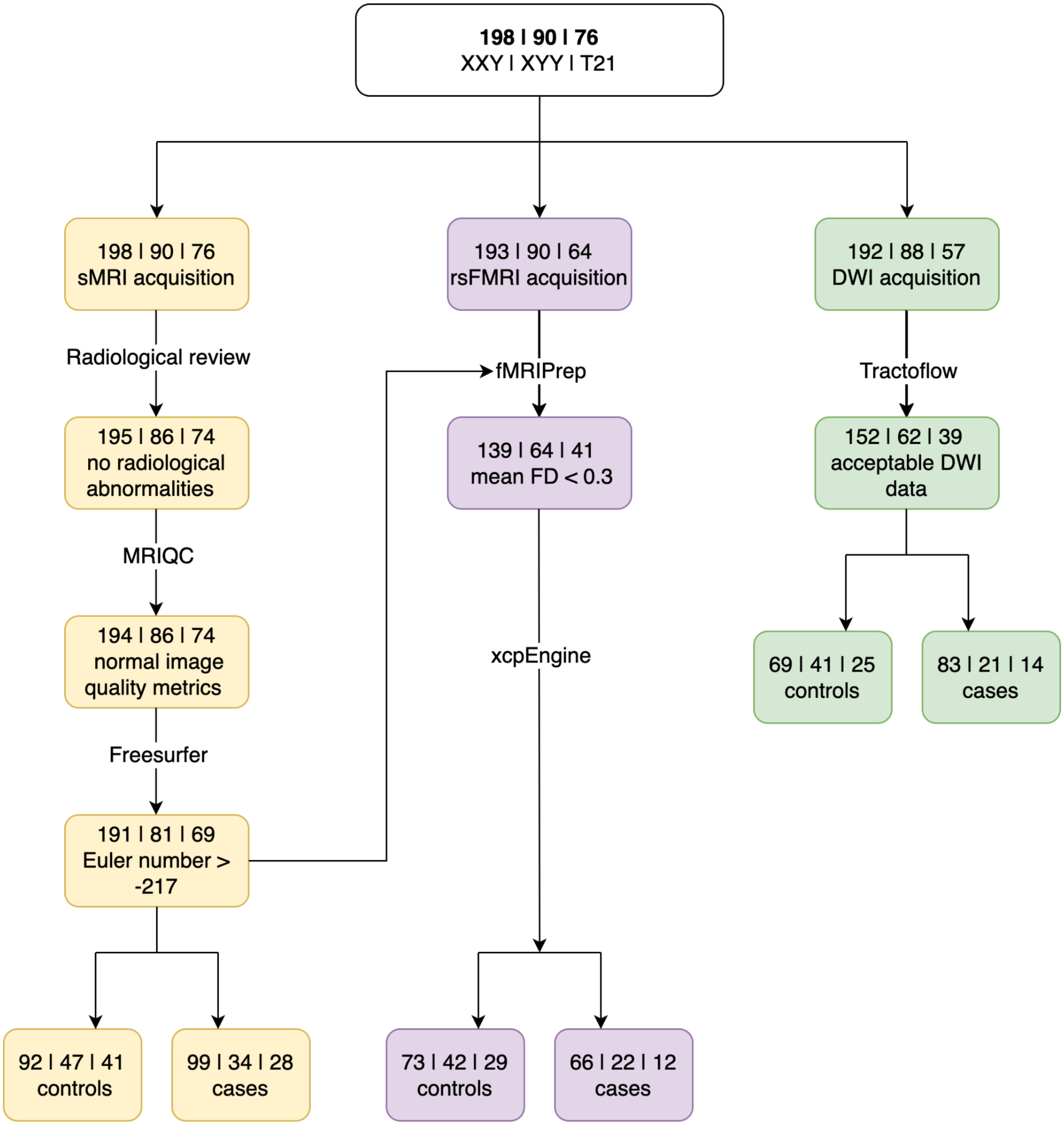


*Figure S1. Quality control workflow. The sMRI workflow is in yellow, rsFMRI in purple, and DWI in green. T1w scans were first assessed by a radiologist for any abnormalities, they were then processed using MRIQC to identify any scans with outlying image quality metrics, and the Freesurfer-generated Euler number was used as a final QC step.*

##### 1.5 Computing significant-intermap similarities

We concatenated the three ROI by ΔIDP matrices into a single 344 ROI by 45 ΔIDP matrix. Given the previously demonstrated impact of spatial auto-correlation on statistical inference when comparing brain maps, we applied a spatial permutation framework to determine which correlations are statistically significant[11–13]. Briefly - we estimated the centroids of the 344 parcels based on their geodesic distance and computed 1000 angular rotations. The ΔIDP maps were rotated using each of the rotations to yield 1000 null ROI by ΔIDP matrices, which were then correlated separately with the original concatenated ROI by ΔIDP matrix. A null distribution of 1000 correlations was generated by storing the largest absolute value (excluding the diagonal) of each permuted correlation matrix, and the null distribution was used to estimate the two-tailed p-values for the correlations in the original RO by ΔIDP matrix.

##### 1.6 Network enrichment

For each aneuploidy, we used the aforementioned 1000 angular rotations to spin the PC1 map, re-average to attain network-specific null distributions, and assess significance by computing a two-tailed p-value between the empirical and null average PC1 values for each network.

##### 1.7 Allen Human Brain Atlas

Regional microarray expression data were obtained from 6 post-mortem brains (1 female, ages 24.0--57.0, 42.50 +/- 13.38) provided by the Allen Human Brain Atlas (AHBA, https://human.brain-map.org)[14]. Data were processed with the abagen toolbox (version 0.1.3; https://github.com/rmarkello/abagen) using a 360-region volumetric atlas in MNI space.

First, microarray probes were reannotated using data provided by ref[15]; probes not matched to a valid Entrez ID were discarded. Next, probes were filtered based on their expression intensity relative to background noise[16], such that probes with intensity less than the background in >=50.00% of samples across donors were discarded. When multiple probes indexed the expression of the same gene, we selected and used the probe with the most consistent pattern of regional variation across donors (i.e., differential stability[17]), calculated with:

$$\Delta_S(p) = \frac{1}{\binom{N}{2}} \sum_{i=1}^{N-1} \sum_{j=i+1}^{N} \rho[B_i(p), B_j(p)]$$

where $\rho$ is Spearman's rank correlation of the expression of a single probe, p, across regions in two donors $B_i$ and $B_j$, and N is the total number of donors. Here, regions correspond to the structural designations provided in the ontology from the AHBA.

The MNI coordinates of tissue samples were updated to those generated via non-linear registration using the Advanced Normalization Tools (ANTs; https://github.com/chrisfilo/alleninf). Samples were assigned to brain regions in the provided atlas if their MNI coordinates were within 2 mm of a given parcel. To reduce the potential for misassignment, sample-to-region matching was constrained by hemisphere and gross structural divisions (i.e., cortex, subcortex/brainstem, and cerebellum, such that e.g., a sample in the left cortex could only be assigned to an atlas parcel in the left cortex)[15]. If a brain region was not assigned a tissue sample based on the above procedure, every voxel in the region was mapped to the nearest tissue sample from the donor in order to generate a dense, interpolated expression map. The average of these expression values was taken across all voxels in the region, weighted by the distance between each voxel and the sample mapped to it, in order to obtain an estimate of the parcellated expression values for the missing region.

Inter-subject variation was addressed by normalizing tissue sample expression values across genes using a robust sigmoid function[18]:

$$x_{norm} = \frac{1}{1 + \exp(-\frac{(x - \langle x \rangle)}{\mathrm{IQR}_x})}$$

where $\langle x \rangle$ is the median and $\mathrm{IQR}_x$ is the normalized interquartile range of the expression of a single tissue sample across genes. Normalized expression values were then rescaled to the unit interval:

$$x_{scaled} = \frac{x_{norm} - \min(x_{norm})}{\max(x_{norm}) - \min(x_{norm})}$$

Gene expression values were then normalized across tissue samples using an identical procedure. All available tissue samples were used in the normalization process regardless of whether they were assigned to a brain region. Tissue samples not matched to a brain region were discarded after normalization. Samples assigned to the same brain region were averaged separately for each donor and then across donors, yielding a regional expression matrix.

##### 1.8 ENIGMA analysis

Meta-analytic maps from the ENIGMA consortium were downloaded from the ENIGMA toolbox (https://doi.org/10.1038/s41592-021-01186-4). Available case-control maps included: a CT map for autism spectrum disorder (ASD), CT and SA maps for pediatric, adolescent, and adult populations with attention deficit hyperactivity disorder (ADHD), CT and SA maps for adolescent and adult populations with bipolar disorder (BD), CT and SA maps for adolescent and adult populations with major depressive disorder (MDD), CT and SA maps in pediatric and adult OCD populations, and CT and SA maps in schizophrenia.

#### 2. Results

##### 2.1 Regional ΔIDP changes in XXY, XYY, and T21 with and without TTV correction

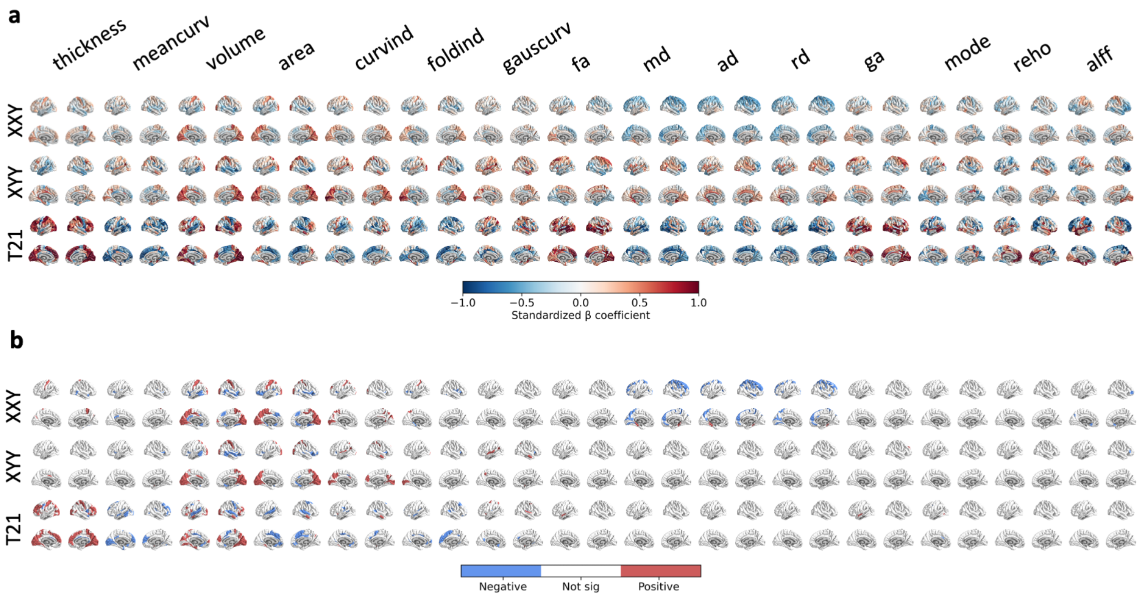


*Figure S2. IDP maps for both hemispheres with TTV correction. a) Unthresholded effect sizes where blue colors denote a decrease in cases relative to controls and red colors denote an increase in cases relative to controls. b) Thresholded effect sizes where all regions of interest colored in red exhibit statistically significant increases in cases relative to controls, and all regions of interest colored in blue exhibit statistically significant decreases in cases relative to controls.*

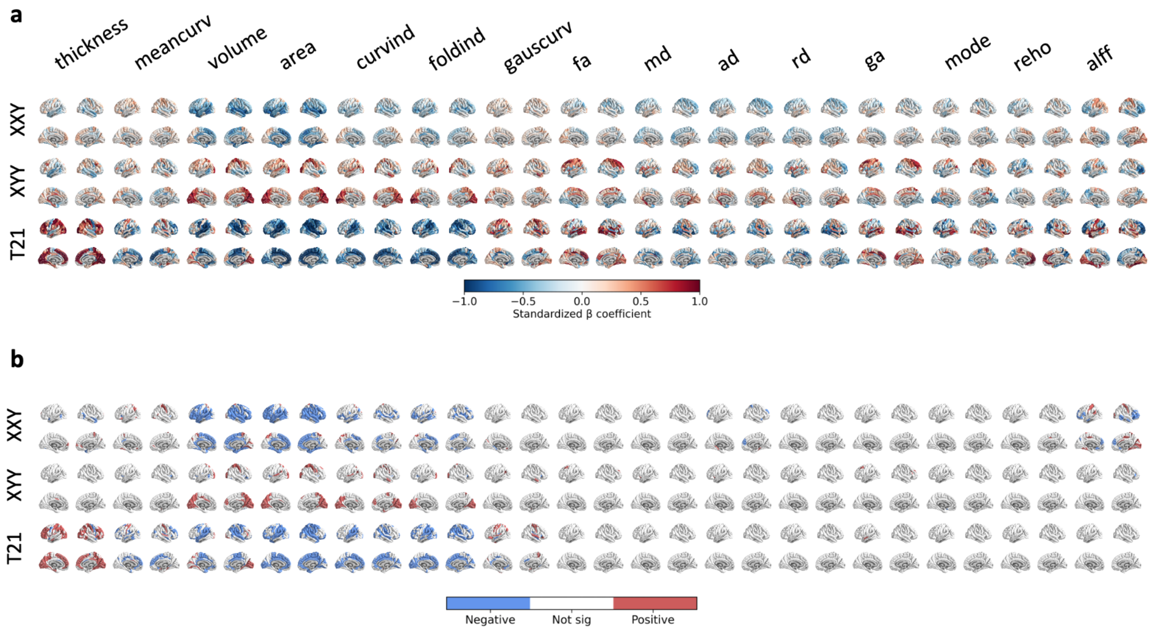


*Figure S3. IDP maps for both hemispheres without TTV correction. a) Unthresholded effect sizes where blue colors denote a decrease in cases relative to controls and red colors denote an increase in cases relative to controls. b) Thresholded effect sizes where all regions of interest colored in red exhibit statistically significant increases in cases*

*relative to controls, and all regions of interest colored in blue exhibit statistically significant decreases in cases relative to controls.*

| | xxy_case | xxy_control | xxy_sig | xyy_case | xyy_control | xyy_sig | t21_case | t21_control | t21_sig |
|---|---|---|---|---|---|---|---|---|---|
| **thickness ($mm$)** | 2.65617 | 2.65868 | | 2.68628 | 2.6997 | | 2.79116 | 2.65652 | True |
| **meancurv** | 0.117099 | 0.116863 | | 0.117079 | 0.116732 | | 0.114144 | 0.11871 | True |
| **volume ($mm^3$)** | 511188 | 549538 | True | 576270 | 551492 | | 457300 | 495690 | True |
| **area ($mm^2$)** | 175319 | 187066 | True | 196347 | 185085 | True | 143209 | 167421 | True |
| **curvind** | 0.787782 | 0.841462 | True | 0.953591 | 0.863248 | True | 0.650945 | 0.777545 | True |
| **foldind** | 8.20155 | 8.78555 | True | 9.55147 | 9.17695 | | 6.22176 | 7.74979 | True |
| **gauscurv** | 0.0251246 | 0.025251 | | 0.0271449 | 0.0258844 | | 0.0258116 | 0.0261298 | |
| **fa** | 0.142403 | 0.142772 | | 0.150532 | 0.147741 | | 0.142758 | 0.138535 | |
| **md** | 0.000941752 | 0.000957731 | | 0.000941154 | 0.0009302 | | 0.000958401 | 0.00096848 | |
| **ad** | 0.00107217 | 0.00108927 | | 0.00107924 | 0.0010652 | | 0.00109251 | 0.00109951 | |
| **rd** | 0.000876542 | 0.00089196 | | 0.000872111 | 0.000862698 | | 0.000891349 | 0.000902968 | |
| **ga** | 0.210705 | 0.210233 | | 0.224903 | 0.221036 | | 0.206544 | 0.200094 | |
| **mode** | 0.192907 | 0.194145 | | 0.192423 | 0.19745 | | 0.209797 | 0.220343 | |
| **reho** | 0.323804 | 0.33832 | | 0.309824 | 0.325968 | | 0.286859 | 0.295813 | |
| **alff** | 0.13339 | 0.141922 | | 0.130371 | 0.153503 | True | 0.130324 | 0.160057 | |
| **TTV ($mm^3$)** | 1.17395e+06 | 1.27269e+06 | True | 1.29992e+06 | 1.25755e+06 | | 1.01291e+06 | 1.15703e+06 | True |
| **FD ($mm$)** | 0.143242 | 0.12327 | True | 0.127285 | 0.11005 | | 0.161708 | 0.0963897 | True |
| **euler_mean_bh** | -25.0303 | -31.5217 | | -44.8529 | -42.7447 | | -24.3571 | -32.8049 | True |
| **age_sMRI ($years$)** | 16.3808 | 16.2445 | | 15.4912 | 13.9681 | | 15.6821 | 16.122 | |
| **age_DWI ($years$)** | 16.2605 | 16.1733 | | 16.1143 | 13.9 | | 17.6357 | 17.136 | |
| **age_rsFMRI ($years$)** | 16.873 | 17.9295 | | 17.1273 | 14.7214 | | 17.15 | 18.5207 | |

*Table S1. Global measures for all IDPs, TTV, FD, and Euler number. For area and volume we display the total value for each measure across the 344 included HCP ROIs averaged across subjects whereas for the remaining phenotypes, we display the average value for each measure across the 344 included HCP ROIs averaged across subjects. FD is computed using subjects who have functional MRI data while TTV and euler_mean_bh are for all subjects. For each of the phenotype-by-aneuploidy comparisons, we indicate whether the global measures are significantly different between the respective controls and cases from that cohort in the 'xxy_sig', 'xyy_sig', and 't21_sig' columns. Euler_mean_bh = Euler number; TTV = total tissue volume; FD = framewise displacement.*

#### 2.2 Organizing-principles of cortical change across IDPs, regions and aneuploidies without TTV correction

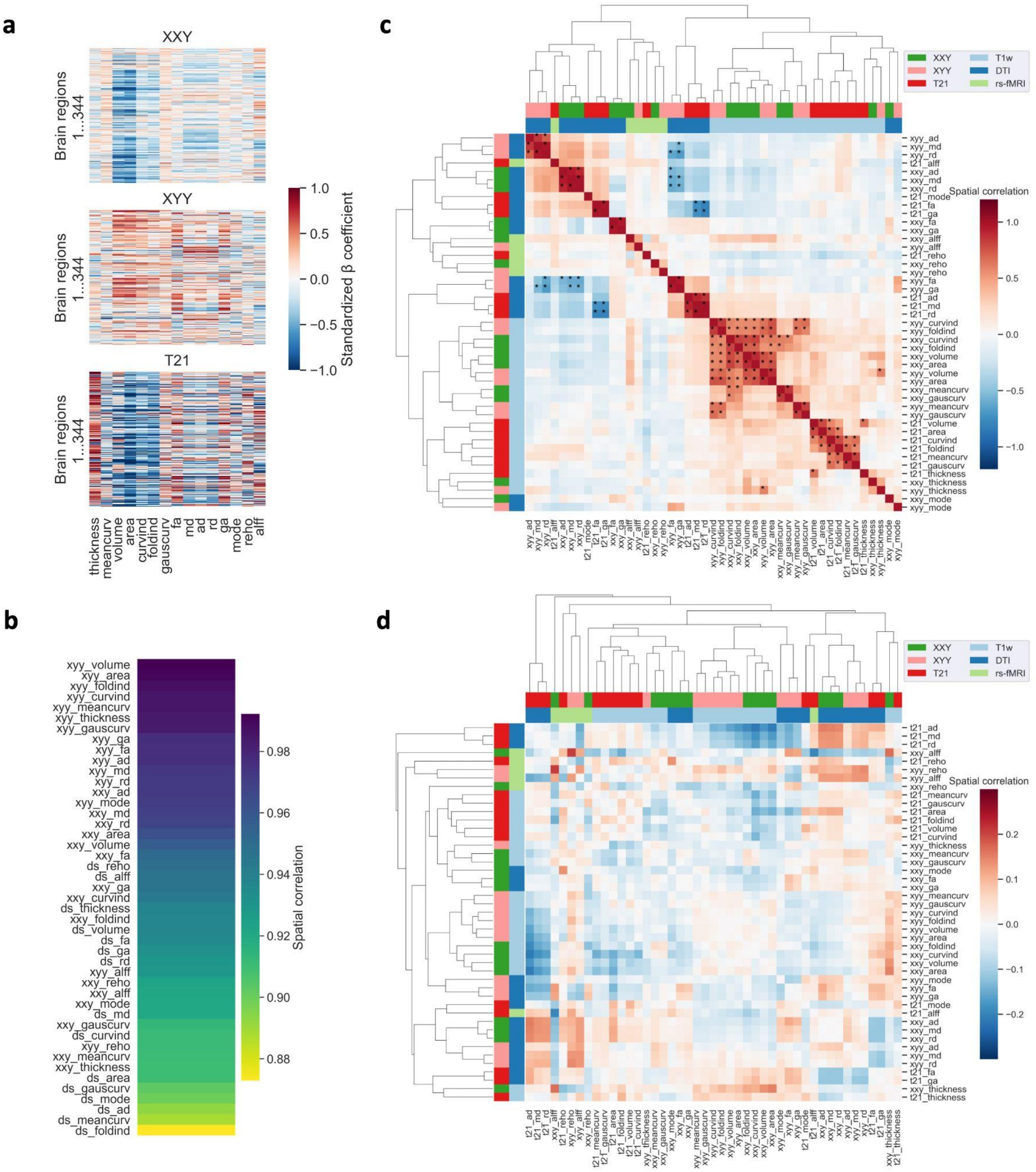


*Figure S4. Visualizing aneuploidy specific ROI by ΔIDP maps and cross-phenotypic relationships without correction for total tissue volume. a) ROI by ΔIDP matrices for each phenotype, where each cell represents the standardized beta coefficient for group effect on a particular aneuploidy at a given brain region. b) Correlation of ROI by ΔIDP maps across all aneuploidies with and without correction for total tissue volume. c) Hierarchical clustering matrix of the concatenated 344 by 45 ROI by ΔIDP matrix. Each row and column corresponds to an aneuploidy-IDP combination. d) Hierarchical clustering matrix of the difference between the cross-correlation matrix generated using the ROI by ΔIDP matrices with TTV correction minus the ROI by ΔIDP matrices without TTV correction.*

#### 2.3 Principal components of multimodal cortical change within and across aneuploidies

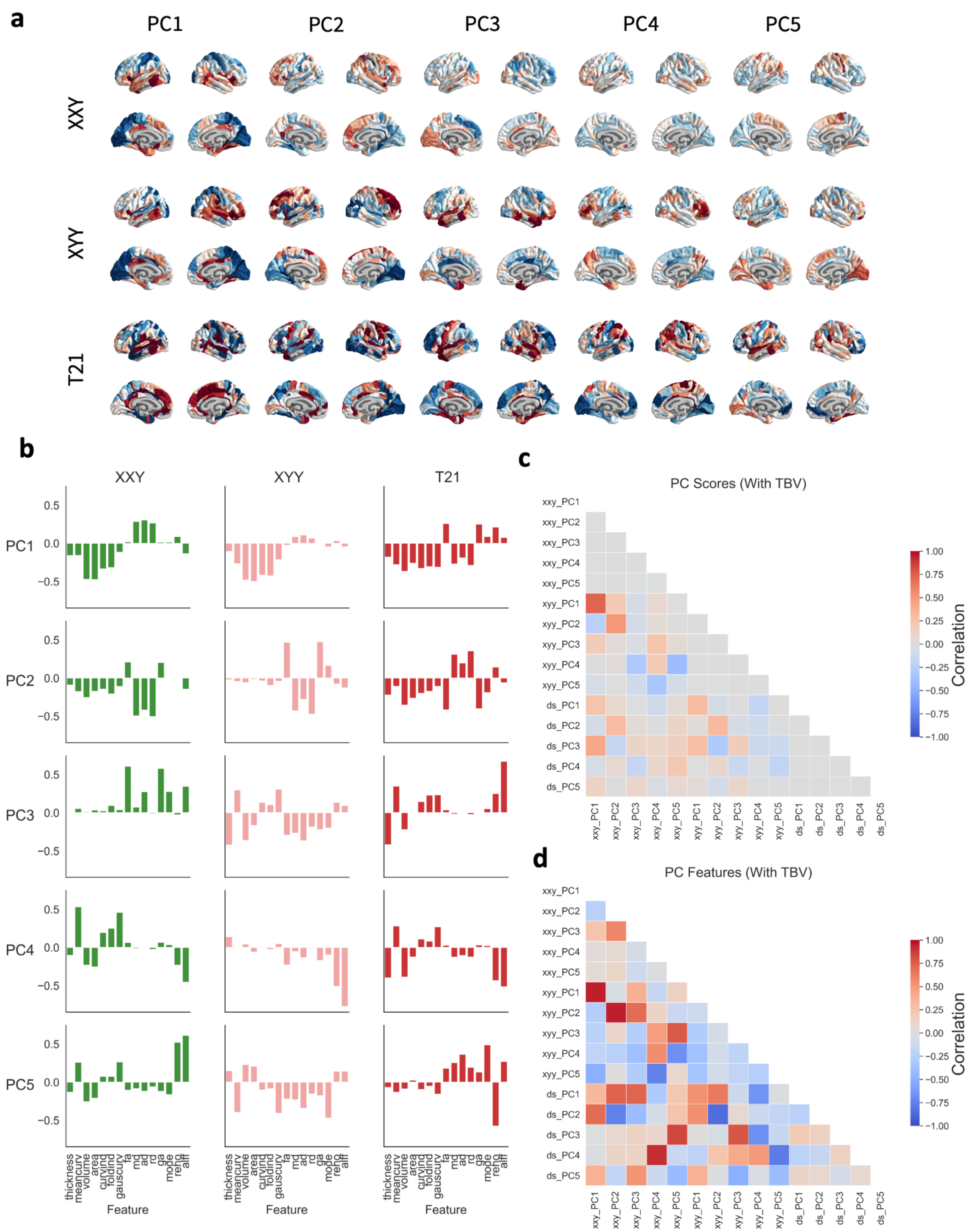

*Figure S5. PCA results for the analytical stream with TTV correction. a) Scores for the first five principal components projected onto the surface for each aneuploidy. b) Feature weights for the first five principal components. c) Cross-correlation of each aneuploidy-PC score combination. d) Cross-correlation of each aneuploidy-PC feature combination.*

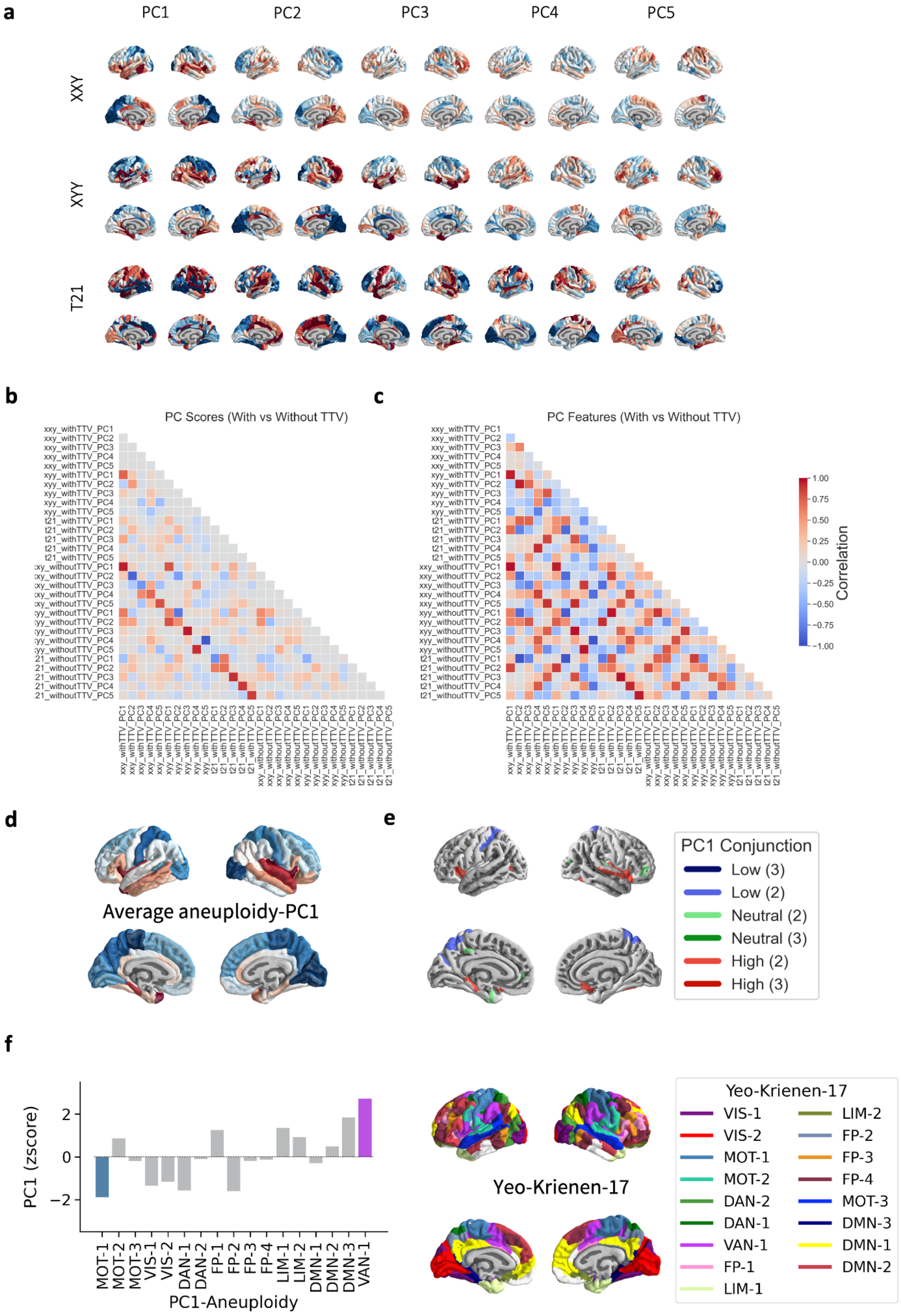
a
PC1
PC2
PC3
PC4
PC5
XXY
XYY
T21
b
PC Scores (With vs Without TTV)
xxy_withTTV_PC1
xxy_withTTV_PC2
xxy_withTTV_PC3
xxy_withTTV_PC4
xxy_withTTV_PC5
xyy_withTTV_PC1
xyy_withTTV_PC2
xyy_withTTV_PC3
xyy_withTTV_PC4
xyy_withTTV_PC5
t21_withTTV_PC1
t21_withTTV_PC2
t21_withTTV_PC3
t21_withTTV_PC4
t21_withTTV_PC5
xxy_withoutTTV_PC1
xxy_withoutTTV_PC2
xxy_withoutTTV_PC3
xxy_withoutTTV_PC4
xxy_withoutTTV_PC5
xyy_withoutTTV_PC1
xyy_withoutTTV_PC2
xyy_withoutTTV_PC3
xyy_withoutTTV_PC4
xyy_withoutTTV_PC5
t21_withoutTTV_PC1
t21_withoutTTV_PC2
t21_withoutTTV_PC3
t21_withoutTTV_PC4
t21_withoutTTV_PC5
c
PC Features (With vs Without TTV)
Correlation
1.00
0.75
0.50
0.25
0.00
−0.25
−0.50
−0.75
−1.00
d
Average aneuploidy-PC1
e
PC1 Conjunction
Low (3)
Low (2)
Neutral (2)
Neutral (3)
High (2)
High (3)
f
PC1 (zscore)
2
0
−2
MOT-1
MOT-2
MOT-3
VIS-1
VIS-2
DAN-1
DAN-2
FP-1
FP-2
FP-3
FP-4
LIM-1
LIM-2
DMN-1
DMN-2
DMN-3
VAN-1
PC1-Aneuploidy
Yeo-Krienen-17
Yeo-Krienen-17
VIS-1
VIS-2
MOT-1
MOT-2
DAN-2
DAN-1
VAN-1
FP-1
LIM-1
LIM-2
FP-2
FP-3
FP-4
MOT-3
DMN-3
DMN-1
DMN-2

*Figure S6. PCA results for the analytical stream without TTV correction. **(a)** Scores for the first five principal components projected onto the surface for each aneuploidy. **(b)** Cross-correlation of each aneuploidy-PC score combination both with and without TTV correction. **(c)** Cross-correlation of each aneuploidy-PC feature combination both with and without TTV correction. **(d)** Average PC1 map (Aneuploidy-PC1) projected onto the cortical surface (in the HCP/Glasser atlas parcellation). (**e**) Conjunction map showing ROIs which sit at the top (red), middle (greens) and bottom (blues) deciles of PC1 scores across aneuploidy/GDD. (**f**) Enrichment of functional connectivity modules defined by Yeo-Krienen-17 (top cortical surfaces, mapped to the Glasser parcellation) for extreme PC1 scores within the Aneuploidy-PC1 map.*

#### 2.2 PC1 annotation

For each aneuploidy-specific PC1 map and the average PC1 map, we first evaluated if the observed omnibus effect of each cortical parcellation on PC1 scores exceeded the distribution of effects from 1000 spatial permutations of the PC1 map. For the Yeo-Krienen-17 parcellation, the average PC1 map shows significant alignment with or without correction for TTV whereas there are more mixed results with the aneuploidy-specific PC1 maps.

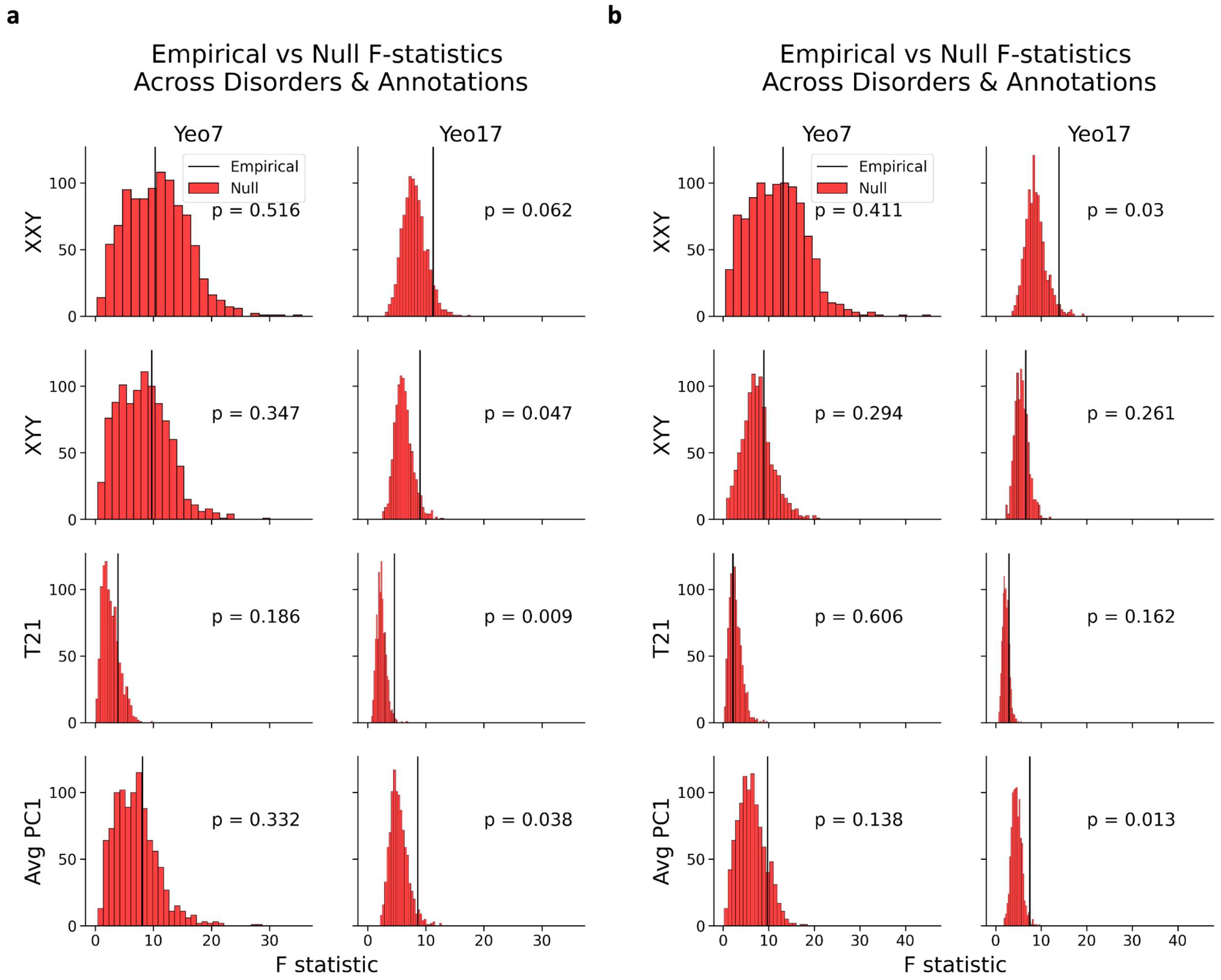


*Figure S7. Empirical versus null F-statistics across aneuploidy PC1 maps and annotations. a) Results using data computed with TTV correction. b) Results using data computed without TTV correction. For both panels, the null distributions of the F-statistics are colored in red, and the empirical F-statistic is in black.*

We assessed enrichment of the individual aneuploidy PC1 maps and found that low PC1 scores were enriched in a somatomotor (MOT-1) subnetwork for XXY, one visual (VIS-2) subnetwork for XYY, and dorsal attention (DAN-2) and default mode (DMN-2) subnetworks for T21 (**Sup Fig 7**). High PC1 scores were enriched in a ventral attention subnetwork (VAN-1) for XYY, and two somatomotor (MOT-2, MOT-3) and one limbic (LIM-2) subnetwork for T21. We observed some changes as well when performing principal component analysis on data that had not been corrected for TTV. In particular, there is disparity with regard to the Yeo-17 functional networks in which each aneuploidy's PC1 score is enriched. Low PC1 scores were enriched in somatomotor (MOT-1) and visual (VIS-2) subnetworks for XXY; dorsal attention (DAN-1), frontoparietal (FP-3), and default mode (DMN-3) subnetworks for XYY; somatomotor (MOT-3), frontoparietal (FP-3), and limbic (LIM-2), and dorsal attention (DAN-2) and default mode (DMN-2) subnetworks for T21. High PC1 scores were enriched in a ventral attention subnetwork (VAN-1) for XYY, and two somatomotor (MOT-2, MOT-3) and one limbic (LIM-2) subnetwork for T21.

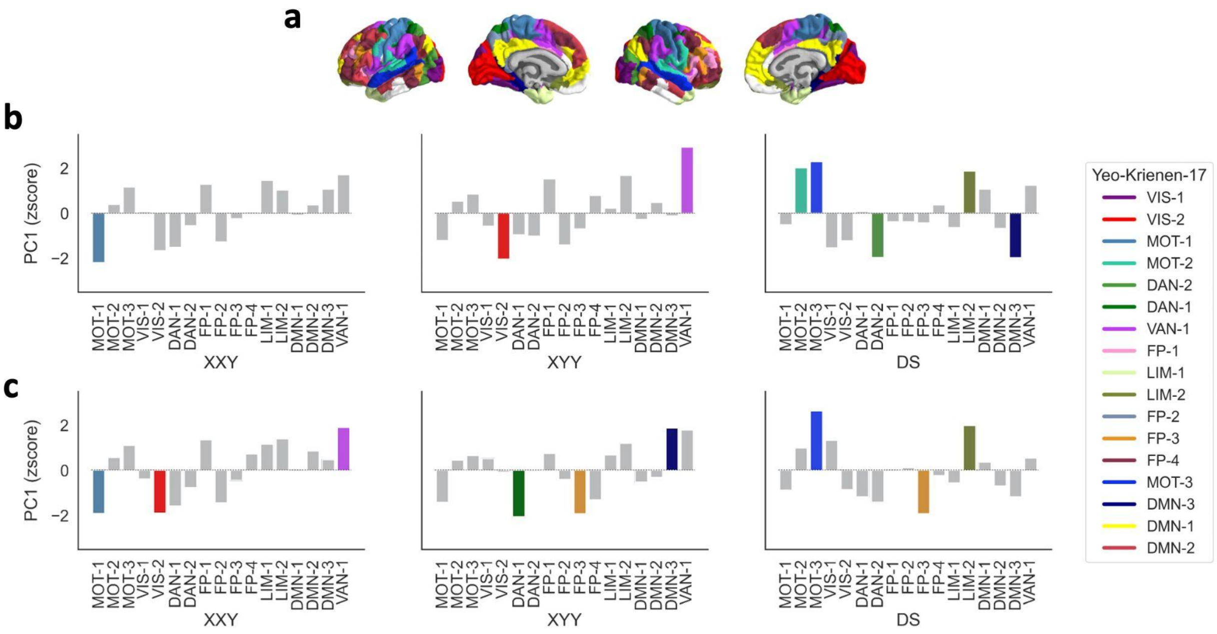


*Figure S8. Enrichment of functional connectivity modules defined by Yeo-Krienen-17 for extreme PC1 scores within XXY, XYY, and T21 PC1 maps. a) Cortical surfaces, mapped to the Glasser parcellation. b) Results using PC1 outputs generated using TTV corrected data. c) Results using PC1 outputs generated using TTV uncorrected data.*

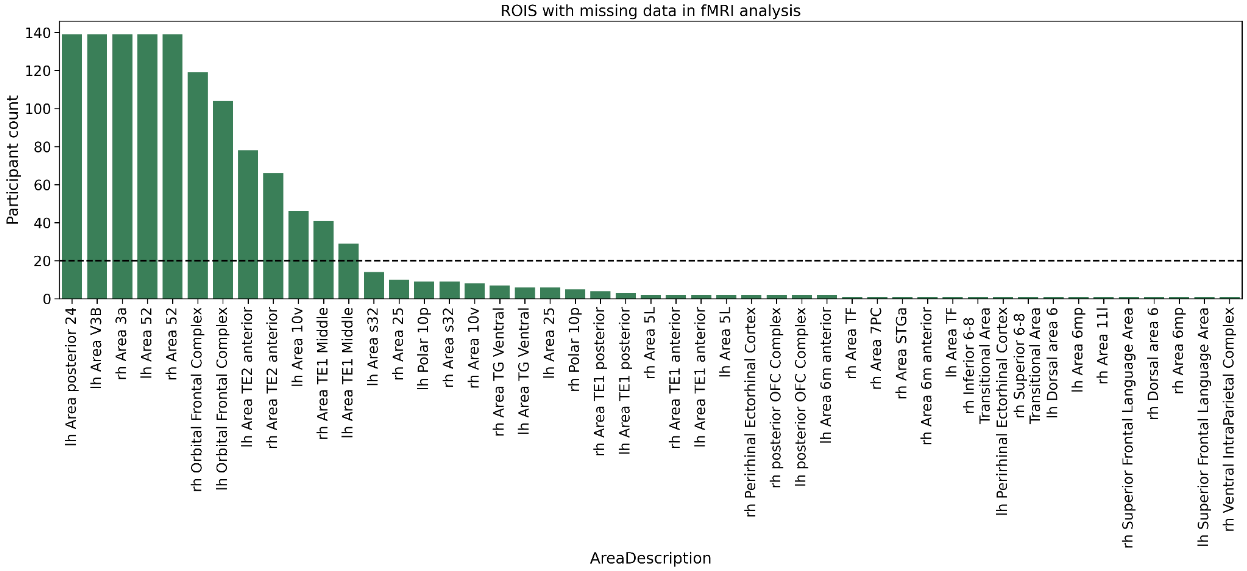


*Figure S9. rs-FMRI signal loss. Using the XXY rsfMRI data, we show the number of participants who exhibit signal loss for particular regions of interest following processing with xcpEngine. ROIs lacking coverage for at least 20 participants were bilaterally excluded from the analysis for each aneuploidy.*

#### References


1. Tustison NJ, Avants BB, Cook PA, et al. N4ITK: Improved N3 Bias Correction. *IEEE Trans Med Imaging*. 2010;29(6):1310-1320. doi:10.1109/TMI.2010.2046908
2. Avants B, Epstein C, Grossman M, Gee J. Symmetric diffeomorphic image registration with cross-correlation: Evaluating automated labeling of elderly and neurodegenerative brain. *Med Image Anal*. 2008;12(1):26-41. doi:10.1016/j.media.2007.06.004
3. Zhang Y, Brady M, Smith S. Segmentation of brain MR images through a hidden Markov random field model and the expectation-maximization algorithm. *IEEE Trans Med Imaging*. 2001;20(1):45-57. doi:10.1109/42.906424
4. Reuter M, Rosas HD, Fischl B. Highly accurate inverse consistent registration: A robust approach. *NeuroImage*. 2010;53(4):1181-1196. doi:10.1016/j.neuroimage.2010.07.020
5. Klein A, Ghosh SS, Bao FS, et al. Mindboggling morphometry of human brains. Schneidman D, ed. *PLOS Comput Biol*. 2017;13(2):e1005350. doi:10.1371/journal.pcbi.1005350
6. Fonov V, Evans A, McKinstry R, Almli C, Collins D. Unbiased nonlinear average age-appropriate brain templates from birth to adulthood. *NeuroImage*. 2009;47:S102. doi:10.1016/S1053-8119(09)70884-5
7. Wang S, Peterson DJ, Gatenby JC, Li W, Grabowski TJ, Madhyastha TM. Evaluation of Field Map and Nonlinear Registration Methods for Correction of Susceptibility Artifacts in Diffusion MRI. *Front Neuroinformatics*. 2017;11. doi:10.3389/fninf.2017.00017
8. Huntenburg J, Str L. Evaluating nonlinear coregistration of BOLD EPI and T1w images. :29.
9. Ciric R, Rosen AFG, Erus G, et al. Mitigating head motion artifact in functional connectivity MRI. *Nat Protoc*. 2018;13(12):2801-2826. doi:10.1038/s41596-018-0065-y
10. Behzadi Y, Restom K, Liau J, Liu TT. A component based noise correction method (CompCor) for BOLD and perfusion based fMRI. *NeuroImage*. 2007;37(1):90-101. doi:10.1016/j.neuroimage.2007.04.042
11. Alexander-Bloch A, Giedd JN, Bullmore E. Imaging structural co-variance between human brain regions. *Nat Rev Neurosci*. 2013;14(5):322-336. doi:10.1038/nrn3465
12. Váša F, Seidlitz J, Romero-Garcia R, et al. Adolescent Tuning of Association Cortex in Human Structural Brain Networks. *Cereb Cortex*. 2018;28(1):281-294. doi:10.1093/cercor/bhx249
13. Markello RD, Misic B. *Comparing Spatially-Constrained Null Models for Parcellated Brain Maps*. Neuroscience; 2020. doi:10.1101/2020.08.13.249797
14. Hawrylycz MJ, Lein ES, Guillozet-Bongaarts AL, et al. An anatomically comprehensive atlas of the adult human brain transcriptome. *Nature*. 2012;489(7416):391-399. doi:10.1038/nature11405
15. Arnatkevičiūtė A, Fulcher BD, Fornito A. A practical guide to linking brain-wide gene expression and neuroimaging data. *NeuroImage*. 2019;189:353-367. doi:10.1016/j.neuroimage.2019.01.011
16. Quackenbush J. Microarray data normalization and transformation. *Nat Genet*. 2002;32(S4):496-501. doi:10.1038/ng1032
17. Hawrylycz M, Miller JA, Menon V, et al. Canonical genetic signatures of the adult human brain. *Nat Neurosci*. 2015;18(12):1832-1844. doi:10.1038/nn.4171
18. Fulcher BD, Little MA, Jones NS. Highly comparative time-series analysis: the empirical structure of time series and their methods. *J R Soc Interface*. 2013;10(83):20130048. doi:10.1098/rsif.2013.0048
